# Genetic mining of newly isolated Salmophages for phage therapy

**DOI:** 10.1101/2022.06.14.493971

**Authors:** J. Gendre, M. Ansaldi, D.R. Olivenza, Y. Denis, J. Casadesús, N. Ginet

## Abstract

*Salmonella enterica* - a Gram negative zoonotic bacterium – is mainly a food-borne pathogen and the main cause of diarrhea in humans worldwide. Main reservoirs are found in poultry farms but also in wild birds. The development of antibiotic resistance in *S. enterica* species raises concerns about the future of efficient therapies against this pathogen and revives the interest in bacteriophages as a useful therapy against bacterial infections. Here we aimed at deciphering and functionally annotate 10 new *Salmonella* phage genomes isolated in Spain in the light of phage therapy. We designed a bioinformatic pipeline using available building blocks to *de novo* assemble genomes and perform syntaxic annotation. We then used genome-wide analyses for taxonomic annotation enabled by vContact2 and VICTOR. We were also particularly interested in improving functional annotation using remote homologies detection and comparisons with the recently published phage-specific PHROG protein database. We finally searched for useful functions for phage therapy such as systems encoded by the phage to circumvent cellular defenses with a particular focus on anti-CRIPSR proteins. We thus were able to genetically characterized nine virulent phages and one temperate phage and identified putative functions relevant to the formulation of phage cocktails for *Salmonella* biocontrol.

## II. INTRODUCTION

The massive and often indiscriminate usage of antibiotics for decades in human and animal healthcare as well as in agriculture gives rise to antibiotic multi-resistance in bacterial pathogens; this has become a major concern worldwide as it leads to an increasing number of therapeutic failures. According to current projections reported by the World Health Organization (WHO), infectious diseases could once again become one of the leading causes of death in the world by 2050 and cause dramatic damages to the world economy [1]. The “One Health” concept recognizes the interdependency between the environment, human health and animal health, it is promoted by various international organizations such as the WHO, the Organization for Animal Health (OIE) and the Food and Agriculture Organization (FAO) (see for a definition the FAO website at https://www.fao.org/one-health/en). One Health aims, among other objectives, at promoting the rational use of antibiotics and the development of alternative strategies to tackle bacterial infections.

Bacteriophages (or phages) are viruses that infect bacteria and thus represent as anti-bacterial agents one such promising alternative (or complement) to antibiotics. A given bacteriophage specifically infects a more or less narrow range of bacteria, sometimes down to the species or serotype level. Furthermore, bacteriophages are very diverse, generally easy to isolate and produce. Bacteriophages were independently discovered during the First World War by Frederick W. Twort in 1915 [2] and by Félix d’Hérelle in 1917 [3] who coined the term “bacteriophage” (see Félix d’Hérelle seeding article presented in English by Dr. Roux [4]). Very rapidly in 1918 at a time when antibiotics didn’t even exist, Félix d’Hérelle foresaw how bacteriophages could be used against bacterial infections and successfully utilized them to treat dysenteria [5]. Phage therapy was born and developed to treat both humans and animals. However, in the 1940s phage therapy was supplanted by the emergence of antibiotics and was abandoned in the Western world while it was kept vivid until today in the former Eastern countries and former USSR member states [6]. Nowadays there is a renewed interest worldwide for bacteriophages and phage therapy in the search for an alternative to antibiotics to tackle multi-resistant bacteria and develop targeted antibacterial therapies.

*Salmonella enterica* is a Gram-negative zoonotic bacterium and a major cause of diarrhea worldwide. Avian breeding farms are the main reservoirs for the *Salmonella* species, transmitted to humans by food, especially by eggs. In the concept of “One Health”, treating these reservoirs would have a positive impact on human health. Isolation and characterization of phages targeting *Salmonella* is thus important to develop *S. enterica* biocontrol strategies and prevent massive antibiotic resistance in animal reservoirs. Phage therapy consensually relies on the formulation of a cocktail regrouping several different phages in order to avoid the emergence of bacterial resistance – especially cross-resistance - and eventually target a range of bacterial serotypes as broad as it can be. Another beneficial property of bacteriophage cocktails is that they can be designed to target specific pathogens, leaving the rest of the microbiota unaffected, as is not the case with antibiotics and their indiscriminate action against a wide range of bacteria. A recent and relevant study of Nale et *al.* highlights the synergistic effects of a three-phages cocktail against prevalent *Salmonella* serotypes on poultry and pigs [7]. Using *Galleria mellonella* as a model that correlates to large scale animal models, the authors evidenced that in combination their three phages can lyse about 99,97% of the 22 serotypes they investigated. They also showed *in vivo* in the *Galleria mellonella* model that their cocktail was very efficient as a prophylactic agent. Closer to human health issues is the recent report of the successful treatment of a patient infected by un multi-drug resistant *Mycobacterium abscessus* strain by a patient-tailored three-phage cocktails [8]. However, the authors cautiously warn that a generalized approach is far from being a reality.

Just like antibiotherapy, phage therapy collides with bacterial resistance to phage infection and its success relies on the evaluation of such a probability, implying that we should definitively not reproduce the errors we made with antibiotics. Indeed, co-evolution of bacteria and their viruses for eons led to adaptation and acquisition of a high diversity of bacterial anti-phage strategies. The “pan-immune system” of bacteria comprising anti-phage defenses has been nicely reviewed by Berheim and Sorek [9]. These strategies evolved by bacteria can be as simple as selecting for mutations in the phage receptor, preventing phage adsorption by modifying and adapting its lipopolysaccharide (LPS). Of more concern are mutations in non-receptor host factors that can lead to phage cross-resistance as was shown recently in *S. enterica* serovar Typhimurium [10]. Bacteria can also resort to elegant, more complex and specialized defense systems once the phage genome has been injected such as Restriction- Modification (RM) [11], CRISPR-Cas [12], abortive infection (Abi) [13] comprising the recently described cyclic-oligonucleotide-based anti-phage signaling system (CBASS) [14] or bacteriophage Exclusion (BREX) [15–17] systems to name but a few . These systems are often passed on between bacteria via horizontal gene transfers. As we keep discovering new anti-phage defense systems, the selection of phages and the formulation of a cocktail should take bacterial defense against phage infection into consideration. Undoubtedly, co-evolution works both ways, and phages therefore evolved their own arsenal to counter cellular defenses. One can cite anti-RM or anti-CRISPR proteins [18] as well as the newly discovered anti-CBASS and anti-Pycsar proteins [19, 20]. Given the diversity of anti-phage defenses systems, a matching diversity of yet unknown anti-cellular defenses functions is probably buried into the phage genomic “dark-matter” awaiting discovery. The phage “dark-matter” encompasses all those predicted genes in phage genomes and metaviromes with unknown functions [21]. It is of importance to collect as many information as possible on the phage genomes to select the best candidates when foreseeing phage therapy and keep a balance between killing efficiency, susceptibility against cellular defenses and potential antagonistic interactions between the therapeutical phages. A thorough genomic characterization of phage genomes and especially functional annotation are thus of crucial importance for the successful deployment of phage-based biocontrol strategies.

In this study we sequenced and analyzed 10 new bacteriophage genomes infecting *S. enterica* subsp. *enterica* serovar Typhimurium strain ATCC 14028S. These phages were isolated from wastewater and fresh water pounds in the Sevilla area in Spain. Most phages originate from a study by Olivenza et *al.* [22] aiming at developing epigenetic phage biosensors used to identify and select phages from different natural environments, others belong to Dr. M. Ansaldi and Prof. J. Casadesús personal collections. To improve phage gene detection and annotation, we benchmarked five different gene callers for the syntaxic annotation. We could ascribe a taxonomic affiliation for each phage at the genus level, thanks to genome-wide comparative analyses enabled by vContact2 [23] (based on gene-sharing network) and VICTOR (based on genome-wide phylogeny) [24]. For functional annotation of the predicted ORFs, we combined complementary approaches based on remote homologies detection using Hidden Markov Model (HMM) protein profiles comparisons with PHROG [25], a newly published database of viral protein clusters, and the pdb70 database at the Protein Data Bank [26]. Thirdly, we specifically searched for anti-CRISPR proteins (Acr) using the integrated online platform AcrHub [27] relying on machine learning for Acr prediction. Acr are indeed valuable assets when foreseeing phage therapy. We further mined our annotated genomes for functions that may favor or on the contrary disfavor the selection of a phage for phage therapy.

The 10 new phages studied here all belong to the *Caudoviricetes* and are spread among four genera (*Kuttervirus*, *Chivirus*, *Jerseyvirus* and *Lederbervirus*) from three families (*Ackermannviridae*, *Siphoviridae* and *Podoviridae*) with a majority of six *Kuttervirus* phages isolated (*Ackermanviridae*). One phage, *Salmonella* phage Salfasec_13b (*Lederbergvirus*), was predicted as temperate – which is not a desired property for phage therapy. Among the most interesting features we found phage-encoded proteins targeting the cellular defenses against phage infections such as the Restriction-Modification, CRIPSR-Cas and Abortive infection systems. Noteworthy, *Salmonella* phage Se_F3a (*Kuttervirus*) is likely to code for a novel anti-Abi system derived from the bacterial Phage Shock Protein A (PspA), making this phage an interesting candidate for phage therapy with two different anti-Abi systems, one found in all our *Kutterviruses* previously known to allow bacteriophage T4 to resist the Rex Abi system encoded in some prophages and the second found in *Salmonella* phage Se_F3a hypothesized in this study. We incidentally could propose functional annotations for 24 protein families in the PHROG database currently annotated unknown function. Altogether, this work highlights the need to design specific pipelines for phage genomic analyses to unravel the phage genomic “dark matter” and illustrates the importance of prior knowledge of functions encoded in phage genomes for an educated selection of phage candidates for therapeutical cocktails.

## III. MATERIALS AND METHODS

### III.1 Phage lysates and bacterial strain

*Salmonella* phages Se_AO1, Se_EM1, Se_EM2, Se_EM3, Se_EM4, Se_F1, Se_F2, Se_F3, Se_F6, Salfasec_9, Salfasec_10, Salfasec_11 and Salfasec_13 culture lysates were kindly provided by Dr. Olivenza (Departamento de Genética, Facultad de Biologia, Universidad de Sevilla, 41012 Sevilla, Spain). All phages were isolated in the Sevilla area (Spain) from wastewaters or fresh water pounds. Each phage lysate is registered with a unique BioSample accession number under the BioProject accession number PRJNA767534. When required, *Salmonella enterica* subsp. *enterica* serovar Typhimurium ATCC 14028S was used to propagate and titrate bacteriophages. Cells were grown aerobically in either Lysogeny Broth (LB) at 37°C under shaking (180 rpm) for liquid cultures or on LB-1% agar plates incubated at 37°C.

### III.2 Phage propagation, purification and titration

To propagate phages from culture lysates, 200 mL Erlenmeyer flasks containing 25 mL of LB were inoculated with an overnight (ON) culture of ATCC 14028S at an optical density measured at 600 nm (OD_600nm_) of 0.04. Cells were grown until OD_600nm_ reaches 0.4 then inoculated with 100 µL of 0.22 µm-filtered and chloroform-treated culture lysate. The culture was further incubated for about 4 hours. Cells were lysed by the addition of 10% chloroform (vortexed for 15 s and incubated for 5 mn at room temperature, repeated twice). The aqueous supernatant (phage lysate) was recovered after centrifugation (7500 g, 15 min) to pellet the cell debris and separate the aqueous phase from the solvent. The phage lysate was then 0.22 µm-filtered and stored at 4°C. Phage titration by double agar overlay plaque assay was done according to Kropinski et *al.* [28]. Phage titers are expressed in Plaque Forming Unit per mL (PFU/mL). For further phage purification and concentration, all centrifugation steps were done at 4°C, 20 800 g for one hour and buffers used ice-cold. 4 mL of phage lysate (titer between 1.10^10^ and 3.10^11^ PFU/mL) were centrifuged and the pellet resuspended in 2 mL of 0.22 µm-filtered Phage Buffer (PB: 10 mM Tris-HCl pH 7.5, 100 mM NaCl, 25 mM MgCl_2_, 1 mM CaCl_2_). Phages were pelleted once again and resuspended in 200 µL of PB. Purified phage suspensions were stored at 4°C.

### III.3 Phage morphological characterization by TEM

For Transmission Electronic Microscopy (TEM) exploration, the final phage pellet resuspension was performed in 2 mL of 0.22 µm-filtered TEM buffer (0.1 M ammonium acetate pH 7). Phages were pelleted once again and resuspended in 20 µL of TEM buffer. Formvar/carbon coated copper grids were prepared at the Institut de la Méditerranée (IMM) Microscopy facility. 5 µL of the phage solution was pipetted onto the grid surface and allowed to sediment for 3 min at RT. Excess liquid was then blotted and grids negatively stained according to Ackermann’s protocol with 2% uranyl acetate. Observations were made on a FEI Tecnai G2 20 TWIN (200KV), laB6, Gatan Oneview 4kx4k CMOS transmission electron microscope. Images were visualized using FIJI [29].

### III.4 Phage DNA purification

To remove non-encapsidated nucleic acids, 20 µL of DNAse I (6 mg/mL, Euromedex), 2 µL of RNAse A (4 mg/mL, Promega) and 4 µL of DpnI restriction enzyme (20 000 U/mL, NEB) were added to 200 µL of purified phages in PB. The sample was then incubated one hour at 37°C. DNAse I was inactivated by incubating the sample 20 min at 65°C under shaking. Phage DNA was released from the capsid by adding 20 µL of 10% SDS and 20 µL of proteinase K (50 µg/mL, Epicentre) and incubating the sample one hour at 56°C. The sample volume was completed with PB buffer q.s. 600 µL. Phage DNA was extracted by adding 600 µL phenol/chloroform/isoamyl alcohol (25:24:1, Sigma-Aldrich) then vortexing the sample for 30 s. The upper aqueous phase containing the nucleic acids was recovered after centrifugation (10 000 rpm, 10 mn, 4°C). The extraction was repeated twice. The aqueous phase containing DNA was then ethanol-precipitated. The DNA pellet was finally resuspended in 20 µl DNAse/RNase free water and stored at −20°C. Phage dsDNA was quantified with a Qubit™ fluorometer in combination with the Qubit™ dsDNA HS Assay kit (Invitrogen).

Phage genomic DNA purified from *Salmonella* phages Se_F1, Se_F2, Se_F3, Se_F6 and Salfasec_11 phage lysates was mechanically fragmented in 50 µl microtubes using a Covaris M220 sonifier with the following parameters: peak power 75W, duty factor 10%, Cycles/Burst 200.

Phage genomic DNA purified from *Salmonella* phage Se_AO1, Se_ML1, Se_EM1, Se_EM2, Se_EM3, Se_EM4, Salfasec_9, Salfasec_10 and Salfasec_13 phage lysates was enzymatically fragmented by the tagmentase technology from the NEXTERA XT kit (Illumina®) according to the manufacturer’s protocol.

DNA libraries for high throughput sequencing were prepared from the fragmented DNA at the IMM Transcriptomic and Genomic facility with the NEBNext® Ultra™ II DNA Library Prep kit for Illumina® (New England Biolabs) for *Salmonella* phages Se_F1, Se_F2, Se_F3, Se_F6, Salfasec_11 and with the NEXTERA XT kit (Illumina®) for *Salmonella* phage Se_AO1, Se_ML1, Se_EM1, Se_EM2, Se_EM3, Se_EM4 according to the manufacturers’ protocols.

*Salmonella* phages Salfasec_9, Salfasec_10 and Salfasec_13 DNA libraries preparation with the NEXTERA XT kit (Illumina®) was subcontracted to AllGenetics (A Coruña, Spain).

### III.5 High throughput DNA sequencing

Prior to sequencing, *Salmonella* phages Se_AO1, Se_EM1, Se_EM2, Se_EM3, Se_EM4, Se_ML1, Se_F1, Se_F2, Se_F3, Se_F6 and Salfasec_11 DNA libraries were quantified with a Qubit™ dsDNA HS Assay kit (Invitrogen) and their size distribution profiles recorded with the TapeStation 4200 System (Agilent) in combination with the D5000 DNA ScreenTape System (Agilent). Libraries were then diluted at 4 nM in the appropriate buffer. Paired-end (2 x 150 bp) DNA sequencing was performed on the MiSeq sequencer (Illumina®) hosted at the IMM Transcriptomic and Genomic facility with a MiSeq v2 (300-cycles) flow cell (Illumina®) according to the manufacturer’s protocol.

*Salmonella* phages Salfasec_9, Salfasec_10 and Salfasec_13 DNA libraries sequencing was subcontracted to AllGenetics (Coruña, Spain).

Raw sequencing reads (FASTQ files trimmed from their Illumina adaptors) for each BioSample were submitted to the NCBI Sequence Read Archive under the BioProject accession number PRJNA767534. Raw sequencing reads quality was then improved with Trimmomatic [30] using the following parameters: SLIDINGWINDOW:4:25 MINLEN:75 for *Salmonella* phages Se_AO1, Se_ML1, Se_EM1, Se_EM2, Se_EM3 and Se_EM4 or ILLUMINACLIP TruSe3-SE.fasta SLIDINGWINDOW:4:28 MINLEN:75 for all other phages. Only trimmed paired-reads were used for *de novo* genome assembly.

### III.6 Genome de novo assembly

Genome *de novo* assembly was performed with SPAdes 3.14.1 with default parameters. Supplementary Table 1 summarizes the size of the 18 meaningful contig(s) obtained for the 14 BioSamples together with their mean genome coverage. By “meaningful” we mean contigs with significant sequence coverage and length compatible with a phage genome (50 kb average size for dsDNA phages) and contigs that are not contaminants. Contaminants are usually small contigs with low sequencing coverage and whose DNA sequences match with unrelated organisms (Blastn analyses). For most contigs we could identify identical, 77 bp-DNA sequences at each contig extremity due to the SPAdes assembly algorithm. We observed this with circularly permuted genomes. We later removed the right-most copy to obtain the final contigs we considered as circular for downstream syntaxic annotation, although we are well aware that the encapsidated biological DNA molecule is linear. Indeed, we found out that some genes were spanning the contig extremities. Such was the case for *Salmonella* phages Se_F2 and F6b genomes.

### III.7 Prediction of DNA packaging

We used PhageTerm [31] to predict putative phage genome termini as well as the DNA packaging mode. PhageTerm is based on the statistical analysis of the sequence coverage after mapping of the short sequencing reads onto the assembled contig. For a statistically sound result, PhageTerm requires a minimum sequence coverage of 50. Obviously PhageTerm requires that the genome extremities are sequenced and this depends on the technology used to construct the DNA libraries for sequencing. Genome extremities can be recovered when the phage DNA has been mechanically fragmented as described above. This was the case for *Salmonella* phages Se_F1, Se_F2, Se_F6 and Salfasec_11. When the tagmentation has been used to construct the libraries (Nextera XT kit from Illumina®), one cannot recover the extremities of a linear dsDNA, which precludes termini identification by PhageTerm when DNA packaging is initiated by the terminase at fix positions (*cos* or *pac* sites) and ends at a fix position (*cos* site). Tagmentation was used for *Salmonella* phages Se_AO1, Se_ML1, Se_EM1, Se_EM2, Se_EM3, Se_EM4, Salfasec_9, Salfasec_10 and Salfasec_13.

### III.8 Syntaxic annotation

For tRNA prediction we used ARAGORN [32]. For Open Reading Frames (ORF) prediction, we tested 5 different gene callers. Indeed, many gene callers are freely available to predict ORFs from nucleic acid sequences but they rely on different algorithms, leading to variable predictions. We selected four of them optimized for bacteria (AMIGene, Glimmer, MetaGeneAnnotator and Prodigal) and one optimized for phage genomes (Phanotate) [33–37].

Locus tag names were built with the same convention for each phage: an alphanumeric string defining the phage name coupled to an alphanumeric string defining the gene product numbering. For instance, Se_AO1_gp_001 refers to the first ORF predicted for *Salmonella* phage Se_AO1. When present, tRNAs were labelled following the same convention: Se_AO1_tRNA1 refers to the first tRNA predicted for *Salmonella* phage Se_AO1 genome. To ease further genomic comparisons, we decided to start the ORFs numbering with the terminase small subunit-encoding ORF as *gp_001*. All DNA sequences were flipped and/or rotated accordingly.

### III.9 Taxonomic classification

For taxonomic classification of our newly isolated phages, we used two different but complementary genome-wide approaches dedicated to viruses vContact2 and VICTOR. Neither method requires previous knowledge of gene functions. Our aim was to tentatively ascribe a taxonomic affiliation down to the genus level to our newly isolated phages. vContact2 constructs a network describing the relationships between individual genomes and build viral clusters (VC). Each VC comprises viruses sharing a significant number of proteins clusters (PC), each PC regrouping protein orthologs. We used the ProkaryoticViralRefSeq201-Merged database accompanying vContact2 software and a concatenated list of all our phage predicted proteins sequences (FASTA format) to position our newly isolated phages in the viral cluster network after analysis by vContact2. We visualized this network with Cytoscape [38]. In contrast to vContact2, VICTOR (Virus Classification and Tree Building Online Resource) generates phylogenetic trees from sequences (FASTA or GenBank format) supplied by the user (up to 100 genomes). For our phages, we used the amino acid-based classification. Our reference is the current viral taxonomic classification defined by the International Committee on Taxonomy of Viruses (ICTV).

### III.10 Detection of protein sequence remote homologies

We generated Hidden Markov Model (HMM) profiles for each predicted protein sequence of each phage with HHblits from the UniRef30_2020_06 database in order to detect sequence remote homologies [39, 40].

### III.11 PHROG-based functional annotation

For functional annotation, we used the newly published PHROG database dedicated to viral proteins. In this database, viral proteins are distributed among families called “phrog”; each phrog represent a cluster of viral proteins orthologs built using remote homology detection by HMM profile-profile comparisons. At the date of its publication, the PHROG database contains 17,473 (pro)viruses of prokaryotes or archaea for a total of 938,864 proteins. 38,880 clusters were defined and manual inspection led to the annotation of 5,108 phrog clusters, representing 50.6% of the total protein dataset. Following the procedure available on the PHROG website (https://phrogs.lmge.uca.fr/READMORE.php), we used HH-suite [41] to compare each predicted gene product from the phages studied here with the PHROG database and ascribe a phrog and its annotation whenever possible. When several hits were found, we kept the best one for the considered protein. The corresponding phrog number and its functional annotation were then transferred to our newly predicted protein. When no hit with the PHROG database was found or the affiliation to an existing phrog was too distant (probability < 80%, Evalue > 1e-4), the ORF was annotated as a singleton of unknown function according to the PHROG guidelines. The entire list of the best hit for each ORF for each phage before manual curation is supplied in the Supplementary Excel file 3.

### III.12 PDB-based functional annotation

In order to improve the PHROG-based functional annotation for protein still annotated unknown function, we performed with HHblits a comparison of each predicted protein sequence HMM profile with the Protein Data Bank pdb70 database to detect homologies with known protein structures. We kept the first hit for each protein sequence for further consideration when the probability was greater than 80% and the Evalue less than 1e-3. When these criteria were met, we also manually checked the prediction coverage to avoid transferring a PDB annotation based on a predicted structural similarity between a small portion (a domain or a sub-domain) of the PDB hit and the phage protein sequences. After all these considerations, we manually checked for phage ORF ascribed to phrog of unknown function if a reliable structural prediction from the pdb70 could be used instead. If so, a new annotation inferred from the structural prediction was transferred to the corresponding phrog of unknown function (see Table 5). A new annotation for this phrog cluster will be proposed to the PHROG database through a dedicated form on its website (https://phrogs.lmge.uca.fr/suggestions.php).

### III.13 Manual curation

Phage genes sometimes overlap or may even be entirety included in one another. Nevertheless, in some cases these overlaps can reveal misprediction from the syntaxic annotation. Thus, in an attempt to reduce false positive predictions, we manually scanned each genome to detect highly overlapping ORFs. The most common situation we encountered were two highly overlapping ORFs, with one ORF coded on one strand with a strong affiliation to a phrog family while the ORF encoded on the opposite strand had a weak affiliation to a phrog family (probability < 70%, Evalue > 1e-4). If the latter did not show any strong homology with a known structure from the PDB (probability >80%, Evalue < 1e-3), we decided to discard the ORF. As an example of manual curation, for *Salmonella* phage Se_AO1, *gp_014* and *gp_016* almost entirely overlap with *gp_015*, all 3 ORFs predicted on the same strand. According to the phrog affiliation metrics generated by HHblits (see Supplementary Excel file 3), the Gp_015 is strongly affiliated to phrog_519 (Prob. 99.5%, Evalue 1.4e-18, 261 proteins sequences in this phrog family) whereas Gp_014 is weakly affiliated to phrog_32723 (Prob. 55%, Evalue 1.1, 2 proteins sequences in this phrog family) and Gp_016 is weakly affiliated to phrog_8445 (Prob. 23.5%, Evalue 7.4, 13 proteins sequences in this phrog family). In this example we discarded *gp_014* and *gp_016*. In another example such as *gp_212*, *gp_213* and *gp_214*, it was not possible to decide which ORF was significant based on the phrog affiliation metrics: all three ORFs, although overlapping, code for proteins strongly affiliated to a phrog family. In this case, we chose to keep all three ORFs. A last example of manual curation is *gp_222*. Based on phrog affiliation metrics, we classified Gp_222 as a “singleton” of unknown function. But *gp_222* completely overlaps with the predicted reliably predicted tRNA6-Val on the same strand, we thus decided to discard *gp_222*. Table 6 summarizes the final gene content predicted for each phage after manual curation. This procedure was carried out for every genome included in this study.

### III.14 Prediction of anti-CRISPR proteins (Acr)

Until very recently, searching for Acr was tedious, essentially relying on “guilt by association” with anti-CRISPR-associated (Aca) proteins containing “helix-turn-helix” motives. But not all Acr are associated with an Aca. Thanks to the new approaches based on machine learning, it is now possible through to predict Acr found in undescribed genomic environments. AcrHub is a platform that incorporates state-of-the-art Acr predictors and three analytical modules (similarity analysis, phylogenetic analysis and homology network analysis) [27]. The AcrHub database contains 339 experimentally validated and 71,728 predicted Acr proteins. Validated Acr are mostly short proteins (89% between 60 and 160 aa). We ran our phage proteomes in AcrHub on two different Acr prediction algorithms (PaCRISPR [42] and AcRanker [43]). Meaningful score thresholds are over 0.5 for PaCRISPR and over -5 for AcRanker. We then used the AcrHub “Similarity analysis” module to relate our predicted Acr with experimentally validated Acr proteins in the AcrHub database. We excluded overlong proteins (> 200 aa) and those whose query cover was below 40%. When pertinent, we compared the genomic surrounding of our predicted *<u>acr</u>* gene with the one an experimentally validated *acr* gene product.

## IV. RESULTS AND DISCUSSION

In this work, we used 14 phage isolates from wastewater and fresh water pounds from the Sevilla region in Spain, mostly collected by Dr. D. R. Olivenza from the Departamento de Genética (Facultad de Biologia, Universidad de Sevilla, Spain). In particular, *Salmonella* phages Se_AO1, Se_ML1, Se_F1, Se_F2, Se_F3 and Se_F6 were detected and isolated by Olivenza et *al.* using an epigenetic phage biosensor [22]. After phage propagation on *Salmonella enterica* subsp. *enterica* serovar *Typhimurium* strain ATCC 14028S, we concentrated and purified the phages from infected culture lysates then extracted, purified and sequenced their DNA as described in the Materials and Methods section. In total, we obtained 14 sequencing datasets of paired-end, trimmed, FASTQ files.

### IV.1 Phage genome *de novo* assembly

Genome *de novo* assembly with SPAdes led to the identification of a total of 18 meaningful contigs in the entire dataset with contig lengths comprised between 40 and 160 kb, a range compatible with phage genome sizes (Supplementary Table 1). Of note, SPAdes assembled 2 distinct contigs from Salfasec_11, Salfasec_13, Se_F3 and Se_F6 purified DNA, possibly representing two different phages in the same sample. This is in accordance with the TEM images (Figure 1) where we could visualize two different phage morphologies in purified phage suspensions for *Salmonella* phages Salfasec_11 (two siphovirus with different tail lengths), Se_F3 (one siphovirus with a small head and a long tail, one myovirus) and Se_F6 (one siphovirus with a small head and a long tail, one myovirus). In the Salfasec_13 phage suspension we could only identify one myovirus morphology despite the assembly of two distinct genomes with very different sizes (157296 and 59161 bp).

**Figure 1:**
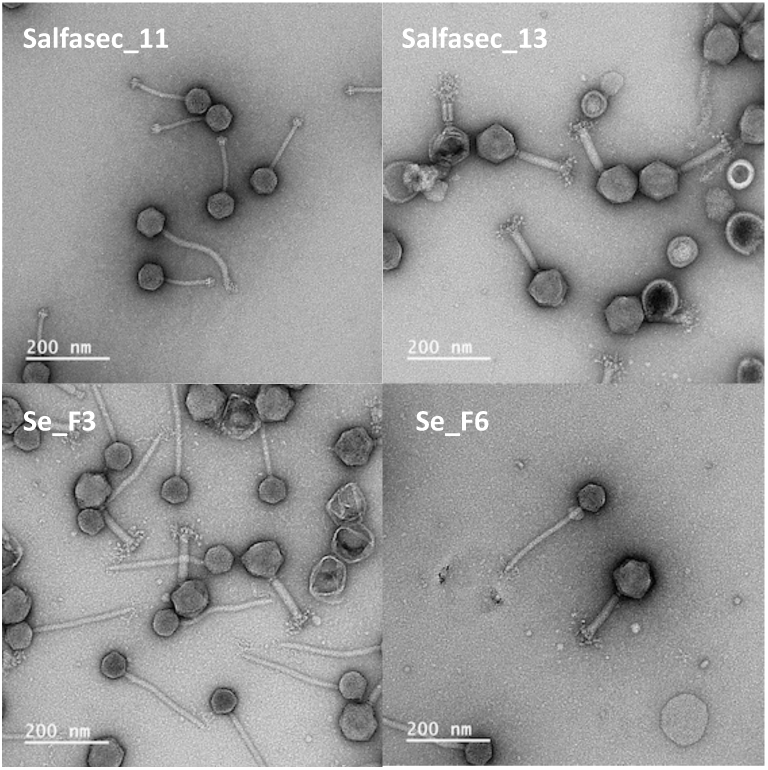
TEM images of negatively stained purified *Salmonella* phages Salfasec_11, Salfasec_13, Se_F3 and Se_F6, suspensions. Phages were purified by centrifugation of 14028S culture lysates and resuspended in TEM buffer. Grids were prepared and visualized as described in the Materials and Methods section. Scale bar: 200 nm.

When relevant, the two DNA sequences present in the same sample were distinguished by adding “a” and “b” to the sample isolate name. Experiences in many labs dealing with phage isolation and sequencing demonstrated that the experimenters are not immune to samples cross-contaminations despite the important precautions taken or that the same phage can be isolated several times from different samples. Resident prophage(s) can also be induced at a basal level in laboratory culture conditions or by the stress triggered by incoming phage infections. In such cases one would expect more than one genome sequenced. As a reminder, our propagation strain *S. enterica* ATCC 14028S is lysogenic for at least three functional temperate bacteriophages, namely Gifsy-1, Gifsy-2 and Gifsy-3 [44], with even a fourth one we predicted as complete and functional using PHASTER [45] *in silico* analysis. We first tested whether the contigs summarized in Supplementary Table 1 represented or not truly different genome sequences. To do so, we computed pairwise DNA sequence similarities - expressed in Average Nucleotide Identity (ANI) - with dRep [46] in order to detect eventual identical species across the samples (Table 1). We chose 99% ANI value as a threshold for identical species.

**Table 1:** Pairwise ANI values calculated with dRep. ANI values ≥ 0.99 (threshold for same species) are highlighted in orange.

| Contig | Salfasec_9 | Salfasec_10 | Salfasec_11a | Salfasec_11b | Salfasec_13a | Salfasec_13b | Se_A01 | Se_EM1 | Se_EM2 | Se_EM3 | Se_EM4 | Se_F1 | Se_F2 | Se_F3a | Se_F3b | Se_F6a | Se_F6b | Se_ML1 |
| --- | --- | --- | --- | --- | --- | --- | --- | --- | --- | --- | --- | --- | --- | --- | --- | --- | --- | --- |
| Salfasec_9 | 1.00 |  |  |  |  |  |  |  |  |  |  |  |  |  |  |  |  |  |
| Salfasec_10 | 1.00 | 1.00 |  |  |  |  |  |  |  |  |  |  |  |  |  |  |  |  |
| Salfasec_11a | 0.00 | 0.00 | 1.00 |  |  |  |  |  |  |  |  |  |  |  |  |  |  |  |
| Salfasec_11b | 1.00 | 1.00 | 0.00 | 1.00 |  |  |  |  |  |  |  |  |  |  |  |  |  |  |
| Salfasec_13a | 0.00 | 0.00 | 0.00 | 0.00 | 1.00 |  |  |  |  |  |  |  |  |  |  |  |  |  |
| Salfasec_13b | 0.00 | 0.00 | 0.00 | 0.00 | 0.00 | 1.00 |  |  |  |  |  |  |  |  |  |  |  |  |
| Se_A01 | 0.00 | 0.00 | 0.00 | 0.00 | 0.94 | 0.00 | 1.00 |  |  |  |  |  |  |  |  |  |  |  |
| Se_EM1 | 0.00 | 0.00 | 0.00 | 0.00 | 0.94 | 0.00 | 0.98 | 1.00 |  |  |  |  |  |  |  |  |  |  |
| Se_EM2 | 0.00 | 0.00 | 0.00 | 0.00 | 0.94 | 0.00 | 0.98 | 0.97 | 1.00 |  |  |  |  |  |  |  |  |  |
| Se_EM3 | 0.00 | 0.00 | 0.00 | 0.00 | 0.94 | 0.00 | 0.98 | 0.99 | 0.98 | 1.00 |  |  |  |  |  |  |  |  |
| Se_EM4 | 0.00 | 0.00 | 0.00 | 0.00 | 0.95 | 0.00 | 0.98 | 0.98 | 0.98 | 0.98 | 1.00 |  |  |  |  |  |  |  |
| Se_F1 | 1.00 | 1.00 | 0.00 | 1.00 | 0.00 | 0.00 | 0.00 | 0.00 | 0.00 | 0.00 | 0.00 | 1.00 |  |  |  |  |  |  |
| Se_F2 | 0.00 | 0.00 | 1.00 | 0.00 | 0.00 | 0.00 | 0.00 | 0.00 | 0.00 | 0.00 | 0.00 | 0.00 | 1.00 |  |  |  |  |  |
| Se_F3a | 0.00 | 0.00 | 0.00 | 0.00 | 1.00 | 0.00 | 0.94 | 0.94 | 0.94 | 0.94 | 0.95 | 0.00 | 0.00 | 1.00 |  |  |  |  |
| Se_F3b | 0.00 | 0.00 | 1.00 | 0.00 | 0.00 | 0.00 | 0.00 | 0.00 | 0.00 | 0.00 | 0.00 | 0.00 | 1.00 | 0.00 | 1.00 |  |  |  |
| Se_F6a | 0.00 | 0.00 | 0.00 | 0.00 | 0.94 | 0.00 | 0.98 | 0.97 | 0.97 | 0.98 | 0.98 | 0.00 | 0.00 | 0.94 | 0.00 | 1.00 |  |  |
| Se_F6b | 0.00 | 0.00 | 0.97 | 0.00 | 0.00 | 0.00 | 0.00 | 0.00 | 0.00 | 0.00 | 0.00 | 0.00 | 0.97 | 0.00 | 0.97 | 0.00 | 1.00 |  |
| Se_ML1 | 0.00 | 0.00 | 0.00 | 0.00 | 1.00 | 0.00 | 0.94 | 0.94 | 0.94 | 0.94 | 0.95 | 0.00 | 0.00 | 1.00 | 0.00 | 0.94 | 0.00 | 1.00 |

According to the values reported in Table 1, we could define four groups of identical DNA sequences: group 1 with *Salmonella* phages Se_EM1 and Se_EM3, group 2 with *Salmonella* phages Se_F1, Salfasec_9, Salfasec_10 and Salfasec_11b, group 3 with *Salmonella* phages Se_F2, Salfasec_11a and Se_F3b and finally group 4 with *Salmonella* phages Se_F3a, Se_ML1 and Salfasec_13a. We could thus reduce our initial dataset from 18 contigs to 10 distinct genomic sequences listed in Table 2. For each group mentioned above, we kept as the group representative only one sequence name. It was either an arbitrary decision for *Salmonella* phage Se_AO1 or when possible motivated by the data we obtained ourselves on our Genomic and Transcriptomic facility (*Salmonella* phages Se_F1, Se_F2 and Se_F3a).

**Table 2:**
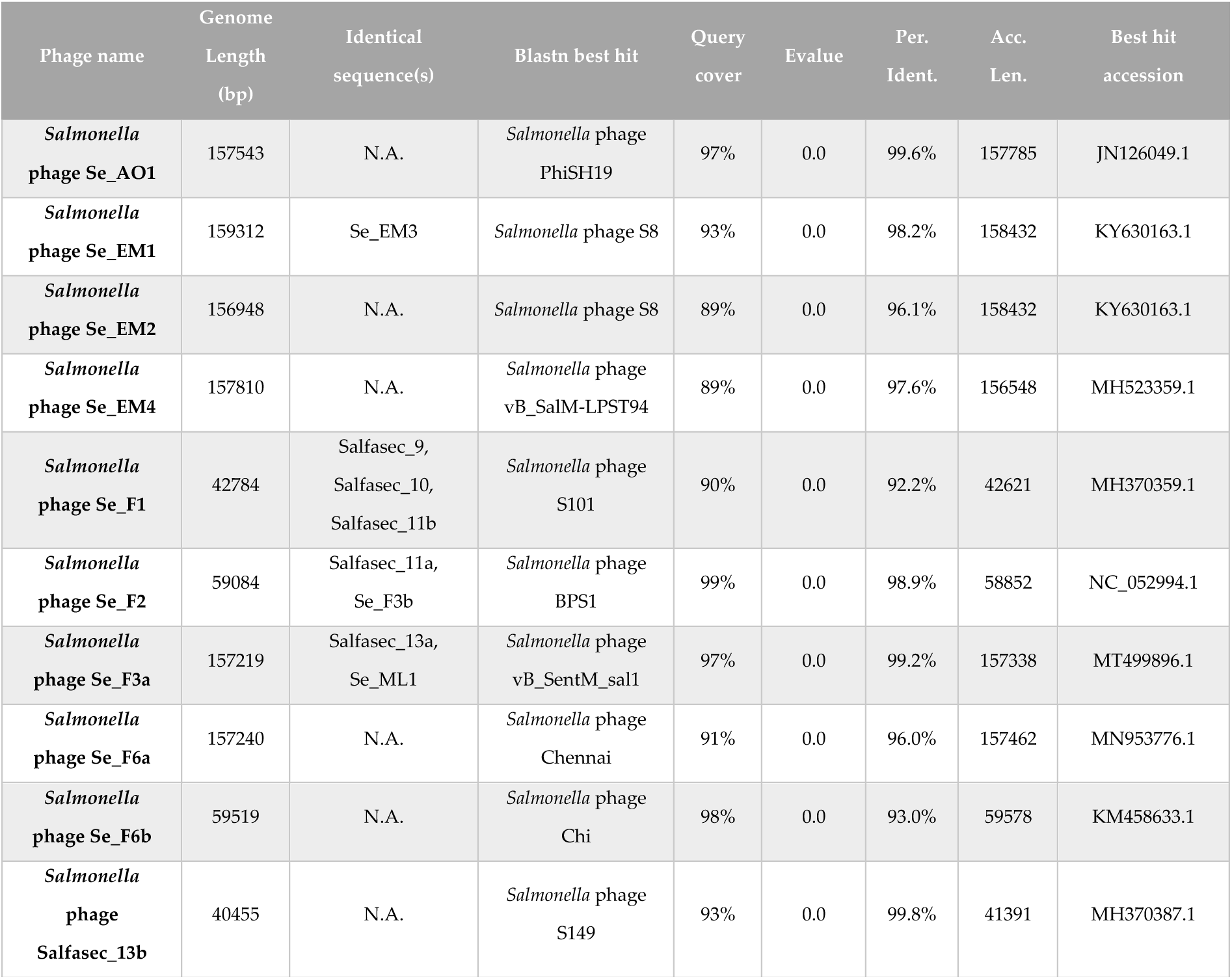
The 10 *Salmonella* phage genomes retained for our downstream genomic analyses. Identical sequences in our samples are indicated by an ANI value ≥ 0.99. We kept for Blastn best-hits only fully sequenced phage genomes (accession numbers in the rightmost column).

| Phage name | Genome Length (bp) | Identical sequence(s) | Blastn best hit | Query cover | Evalue | Per. Ident. | Acc. Len. | Best hit accession |
| --- | --- | --- | --- | --- | --- | --- | --- | --- |
| *Salmonella* phage Se_AO1 | 157543 | N.A. | *Salmonella* phage PhiSH19 | 97% | 0.0 | 99.6% | 157785 | JN126049.1 |
| *Salmonella* phage Se_EM1 | 159312 | Se_EM3 | *Salmonella* phage S8 | 93% | 0.0 | 98.2% | 158432 | KY630163.1 |
| *Salmonella* phage Se_EM2 | 156948 | N.A. | *Salmonella* phage S8 | 89% | 0.0 | 96.1% | 158432 | KY630163.1 |
| *Salmonella* phage Se_EM4 | 157810 | N.A. | *Salmonella* phage vB_SalM-LPST94 | 89% | 0.0 | 97.6% | 156548 | MH523359.1 |
| *Salmonella* phage Se_F1 | 42784 | Salfasec_9, Salfasec_10, Salfasec_11b | *Salmonella* phage S101 | 90% | 0.0 | 92.2% | 42621 | MH370359.1 |
| *Salmonella* phage Se_F2 | 59084 | Salfasec_11a, Se_F3b | *Salmonella* phage BPS1 | 99% | 0.0 | 98.9% | 58852 | NC_052994.1 |
| *Salmonella* phage Se_F3a | 157219 | Salfasec_13a, Se_ML1 | *Salmonella* phage vB_SentM_sal1 | 97% | 0.0 | 99.2% | 157338 | MT499896.1 |
| *Salmonella* phage Se_F6a | 157240 | N.A. | *Salmonella* phage Chennai | 91% | 0.0 | 96.0% | 157462 | MN953776.1 |
| *Salmonella* phage Se_F6b | 59519 | N.A. | *Salmonella* phage Chi | 98% | 0.0 | 93.0% | 59578 | KM458633.1 |
| *Salmonella* phage Salfasec_13b | 40455 | N.A. | *Salmonella* phage S149 | 93% | 0.0 | 99.8% | 41391 | MH370387.1 |

A simple Blastn analysis on the nr/nt database at NCBI (restricted to the Viruses taxon) showed that our 10 distinct DNA sequences are related to known *Salmonella* phages.

Among these, only *Salmonella* phage Salfasec_13b exhibits some small but significant DNA sequence identity with the propagation strain ATCC 14028S genome but only on 3 kb with 89.6% identity. This finding rules out that any of the sequences we obtained originates from resident ATCC 14028S prophages induced during phage propagation.

### IV.2 DNA packaging prediction

When DNA libraries were prepared from randomly fragmented DNA, as was the case for *Salmonella* phages Se_F1, Se_F2, Se_F3a, Se_F6a and Se_F6b, we used PhageTerm to predict putative genome termini and DNA packaging mode. Table 3 summarizes the predictions results.

**Table 3:**
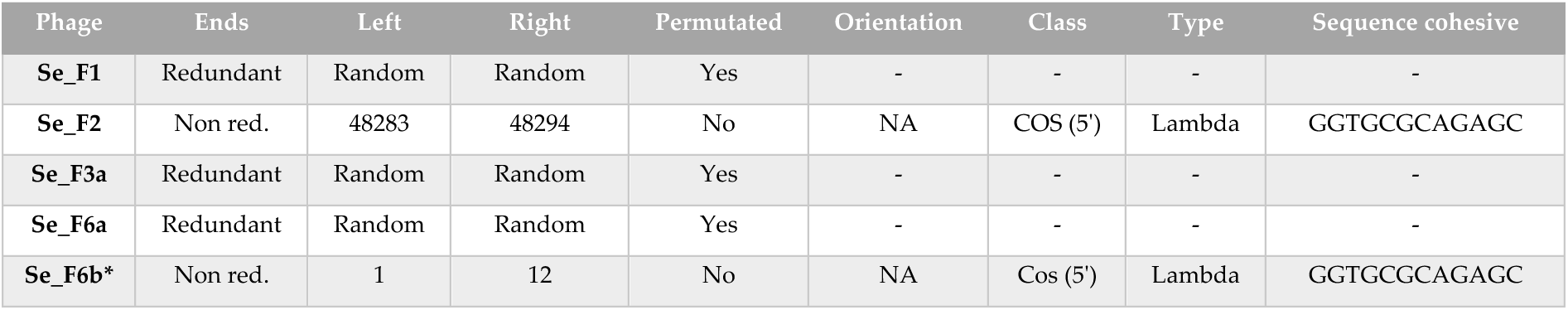
PhageTerm results. *Result to be considered carefully as the mean genome coverage is 15, under the software recommended coverage threshold (50).

| Phage | Ends | Left | Right | Permutated | Orientation | Class | Type | Sequence cohesive |
| --- | --- | --- | --- | --- | --- | --- | --- | --- |
| Se_F1 | Redundant | Random | Random | Yes | - | - | - | - |
| Se_F2 | Non red. | 48283 | 48294 | No | NA | COS (5') | Lambda | GGTGCGCAGAGC |
| Se_F3a | Redundant | Random | Random | Yes | - | - | - | - |
| Se_F6a | Redundant | Random | Random | Yes | - | - | - | - |
| Se_F6b* | Non red. | 1 | 12 | No | NA | Cos (5') | Lambda | GGTGCGCAGAGC |

According to PhageTerm predictions, *Salmonella* phages Se_F1, Se_F3a, Se_F6a use headful (or *pac*) packaging. This is for instance the packaging mode of *Salmonella* phage P22 [47]. This mode of packaging implies that the first cut by the terminase occurs at a *pac* sites on the DNA concatemer issued from the phage DNA replication cycle. The second cut occurs at a random site on the concatemer when the phage capsid is full (more than 1 genome unit length is packaged). The third cut occurs at a random site on the concatemer for the second capsid packaging and so on. Accordingly, at the population level we obtained circularly permutated genomes for *Salmonella* phages Se_F1, Se_F3a, Se_F6a. In contrast, *Salmonella* phages Se_F2 and Se_F6b lied in the *cos* packaging group exemplified by *Escherichia coli* phage Lambda [48], although for *Salmonella* phage Se_F6b the low sequence coverage precludes valid termini prediction by PhageTerm. In this mode of packaging, the concatemer is cut at *cos* sites and each phage capsid contains one identical genome unit length. We could not use PhageTerm for *Salmonella* phages Se_AO1, Se_EM1, Se_EM2, Se_EM4 and Salfasec_13b due to the enzymatic DNA fragmentation method used for the preparation of the DNA libraries that does not allow to recover genome extremities (see Materials and Methods). Nevertheless, we could gain some information on their DNA packaging strategies. Paired-reads mapping with Bowtie2 [49] and mapping visualization with IGV [50] (data not shown) revealed circularly permutated sequences, suggesting that DNA packaging for these phages was probably through headful mechanism.

### IV.3 Syntaxic annotation of 10 newly isolated Salmonella phage genomes

It is common knowledge that predictions vary from one gene caller to another, resulting at the extreme in either missing “real” ORFs or predicting “false” ones, not mentioning discrepancies in initiation codon predictions. ORF prediction is rendered even more complex in bacteriophages due to their genome organization. Indeed, phage genomes are compact, with overlapping genes; in some cases, a gene can be entirely included in another gene, a situation called “overprinting”. This is the case for example for Rz-spanin-o in *E. coli* phage Lambda which is entirely coded in the Rz-spanin-i gene [51]. Phage genomes are often organized in functional modules that can be acquired/modified by recombination with other viruses or mobile genetic elements, or by recombination with their bacterial host and its eventual prophage(s); this mosaic organization resulting from many horizontal gene transfers makes it difficult for any gene caller to build uniform predictive models. The “phage language” is yet to be completely translated and is a current focus of many groups in the field, with great hopes put in Deep Learning techniques.

We thus compared the output of, AMIGene, Glimmer, MetaGeneAnnotator, Phanotate and Prodigal to identify the best algorithm (or combination of algorithms) to optimize gene calling of our phage genomes. We used *Salmonella* phage Se_AO1 as a case study before generalizing the method to the other 9 phage genomes included in this study. In order to compare the different predictions, we built for each gene caller a table where an individual predicted ORF is identified by a stop codon and a coding strand (+ or -) without its predicted initiation codon. This strategy should avoid discrepancies in the predicted initiation codon between the different algorithms. We then compiled the five prediction sets in a single table for each phage with the following unique rule: a predicted ORF from a gene caller is deemed identical to an ORF predicted by another gene caller if they share the same stop codon on the same strand, regardless their respective predicted initiation codon that may be different between the two gene callers. For each phage, we indicated in the table for each predicted ORF whether it is predicted or not by each of the five gene callers. Results are listed in Supplementary Excel file 1 and gene callers’ comparison is illustrated in the Venn diagrams shown in Figure 2 for *Salmonella* phage Se_AO1.

**Figure 2:**
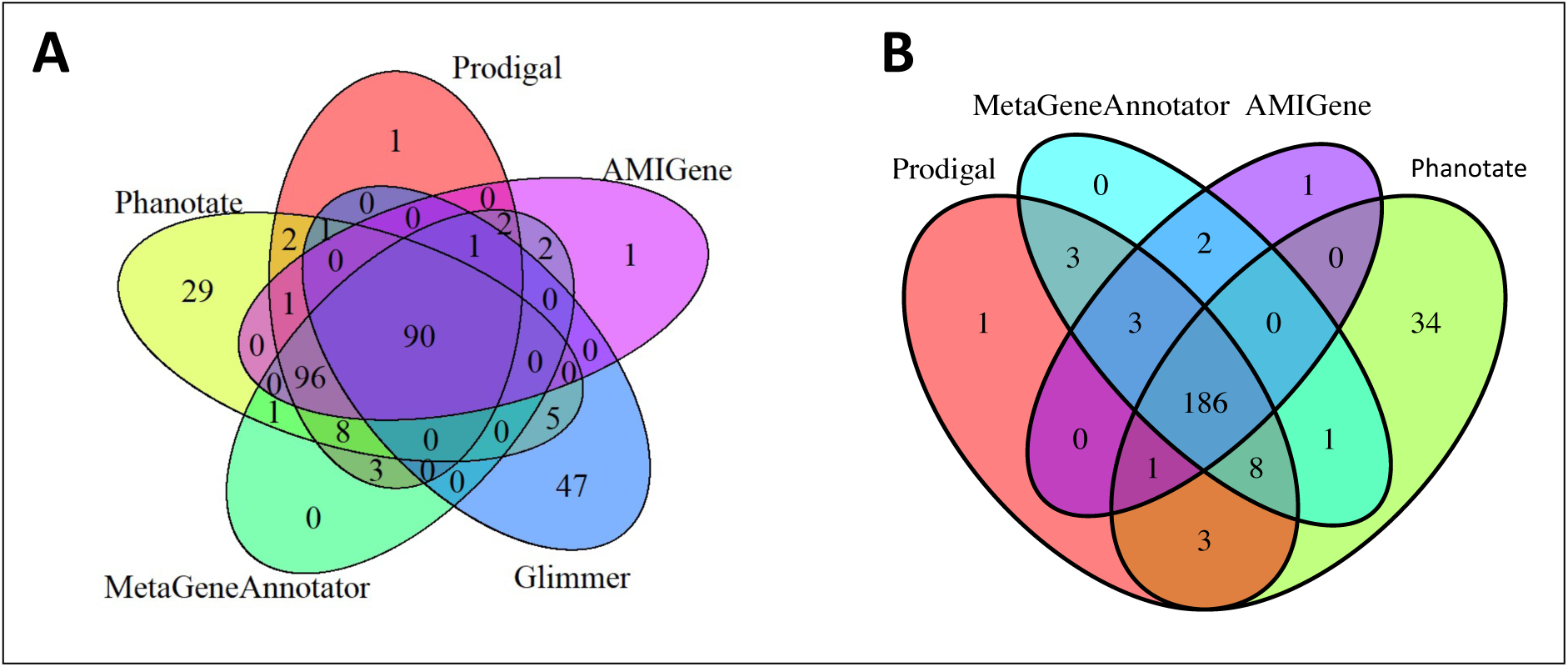
Comparison of the ORFs predictions for *Salmonella* phage Se_AO1 genome. A) Venn diagram for AMIGene, Glimmer, MetaGeneAnnotator, Phanotate and Prodigal predictions (*n* = 290 ORFs). B) Venn diagram omitting Glimmer predictions (*n* = 243 ORFs).

Figure 2A compares the results obtained with all five gene callers. Glimmer displayed the lowest (*n* = 144) and Phanotate the highest (*n* = 233) number of predicted ORFs. A total of *n* = 290 ORFs were predicted, 90 of which were shared by all algorithms (31%). According to this analysis, 69% of the predicted ORFs would depend on the specificity of the gene caller algorithms, which is very improbable. Glimmer predicted *n* = 144 ORFs, 32.6% of which were unique to this software: Glimmer seems then ill-fitted to our purpose. Thus in Figure 2B we drew another Venn diagram omitting Glimmer predictions. On the *n* = 243 predicted ORFs, 76.5% were common to all algorithms (*n* = 186), which is an expected range if all predictors ran with a similar level of accuracy. Of note, Phanotate predicted 34 ORFs overlooked by the three other gene callers. This is not surprising as Phanotate has been purposely designed to identify ORFs in phage genomes.

Setting a cut-off limit for the gene size was not an easy task, as we did not want to keep false predictions or discard ORFs that do code for small proteins. One has to keep in mind that small ORFs in bacteriophages can encode small proteins enabling phages to evade host anti-phage systems. Among them are found: anti-CRISPR proteins (Acr) such as ACR3112-12 (52 aa) from *Pseudomonas* phage D3112 [52]; proteins preventing superinfection such as the immunity protein Imm (83 aa) of bacteriophage T4 [53] or the lipoprotein Llp (77aa) of bacteriophage T5 [54]; anti-RM proteins such as Ocr (117 aa) of bacteriophage T7 [55]; the newly discovered anti-CBASS Acb (about 94 aa) found in an increasing number of phages of various phylogenetic background such as *Pseudomonas* phages PaMx33, 35, 41 and 43 as well as the enterobacteriophage T4 [19, 20].

As the identification of anti-CRISPR and other type of anti-host defenses proteins was one of our goals, we chose a conservative approach as a first approximation and kept all the ORFs predicted by AMIGene, MetaGeneAnnotator, Phanotate and Prodigal for the 10 phage genomes studied here. When several possibilities for the initiation codon were available, we kept the longest version of the ORF. Across our 10 genomes, about 72% of ORFs were predicted by all four gene callers (slightly less for *Salmonella* phage Salfasec_13b) and about 14.5% by Phanotate only (Supplementary Table 2). The detailed ORF predictions for each phage genome are listed in Supplementary Excel file 1.

### IV.4 Taxonomic affiliation

Due to the remarkable diversity of phage nucleotide sequences and the pervasive mosaicism of phage genomes, phage phylogeny does not follow traditional hierarchical phylogeny as noted by Dion et *al.* [56]. Indeed, phage genomes are subjected to fast evolution, thanks to mutations occurring during genome replication, transposition events, recombination events with other genomes as well as imprecise excision during induction. For these reasons, it has always been a challenge to define universal gene markers to classify phages such as the 16S RNA commonly used for bacteria. Depending on the context, good results were obtained with phylogenies based on the capsid major, terminase large subunit or portal protein sequences or a concatenation of some these key structural protein sequences. In the official viral taxonomy maintained by the International Committee on Taxonomy of Viruses taxonomy (ICTV), some taxons were defined by the presence specific genomic features such as a large single subunit RNA polymerase-encoding gene in the *Autographiviridae* family. The current taxonomy of viruses is evolving rapidly using genome-wide analyses, particularly the concept of shared orthologous protein families to infer evolutionary relationships between viral genomes [57, 58]. Indeed, despite a great diversity in phage nucleotide sequences, a remarkable degree of aminoacid sequence homology has been observed as well as structural conservation at the protein level. This is the case for instance for some structural proteins making up the virion particles. Thus, the *Heunggongvirae* kingdom defined by the ICTV is based on the major capsid protein fold first described in bacteriophage HK97 [59] and shared by all *Duplodnaviria*. As described in the Materials and Methods section, we thus resorted to genome wide analyses at the protein level for taxonomic classification with vContact2 and VICTOR. These methods follow the trend in viral taxonomy to replace old classifications methods based on morphologies or phylogenies constructed on a limited set of gene markers. Recently, Turner et *al.* nicely reviewed the challenges of genome-based phage taxonomy and tentatively draw a road map for the future to face them [58]. One important advantage of using such methods to classify phages is that they do not rely on prior knowledge of protein functions. Figure 3A displays a representation generated by Cytoscape of the interaction network between *Caudovirales* bacteriophages generated by vContact2 and including our 10 newly isolated phages. In this representation, the strength of the connection linking two phages depends on the number of proteins clusters they have in common.

**Figure 3:**
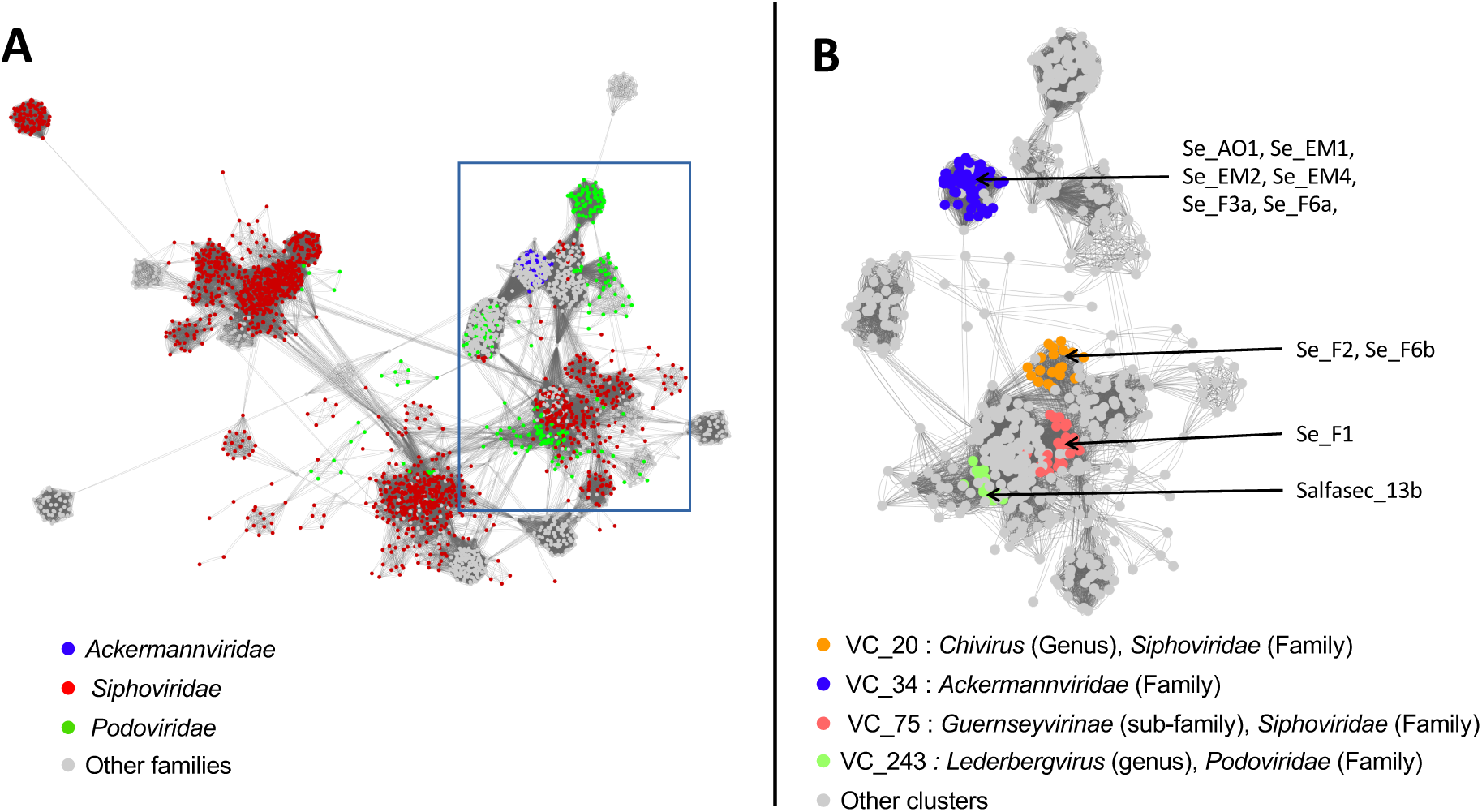
Interaction network between *Caudovirales* bacteriophages based on shared protein ortholog families. The interaction network was built using vContact2 and visualized with Cytoscape. Each dot corresponds to a single phage. Panel A: Network comprising all the *Caudovirales* viruses in the vContact2 database. Viruses belonging to the *Ackermannviridae*, *Siphoviridae* and *Podoviridae* families are highlighted in blue, red and green, respectively. The rectangle indicates the region in the network where the 10 phages studied here are positioned. Panel B: Sub-network of the region underlined in Panel A restricted to *Ackermannviridae*, *Podoviridae* and *Siphoviridae* surrounding our 10 new phages. Viruses belonging to the *Chivirus* genus (VC_20), the *Ackermannviridae* family (VC_34), the *Guernseyvirinae* sub-family (VC_75) and the *Lederbergvirus* genus (VC_243) are highlighted in orange, blue, red and green, respectively. The positions in the sub-network of the 10 new phages are indicated by arrows.

vContact2 also defines viral clusters (VC) regrouping phages that share a significant number of protein clusters. Each VC comprises viruses taxonomically classified at various levels (genus, sub-family or family). Our 10 newly isolated phages fall into four different viral clusters whose members are colored in Figure 3B. *Salmonella* phages Se_F2 and Se_F6b belong to the VC_20 (orange) comprising 22 viruses defined at the genus level (*Chivirus*), *Salmonella* phages Se_AO1, Se_EM1, Se_EM2, Se_EM4, Se_F3a and Se_F6a to the VC_34 (blue) comprising 63 viruses defined at the family level (*Ackermannviridae*), *Salmonella* phage Se_F1 to the VC_75 (red) comprising 25 viruses defined at the sub-family level (*Guernseyvirinae*) and finally *Salmonella* phage Salfasec_13b to the VC_243 (green) comprising 16 viruses defined at the genus level (*Lederbergvirus*). The complete list of the VC generated by vContact2 is available in Supplementary Excel file 2.

For the phages that remained unclassified at the genus level after the previous analysis, we refined the classification with VICTOR using the “amino acid” option analysis to build phylogenetic trees for *Salmonella* phage Se_F1 (Figure 4A) and *Salmonella* phages Se_AO1, Se_EM1, Se_EM2, Se_EM4, Se_F3a and Se_F6a (Figure 4B). In each case, a minimum of three genomes belonging to different families were used as outgroups.

**Figure 4:**
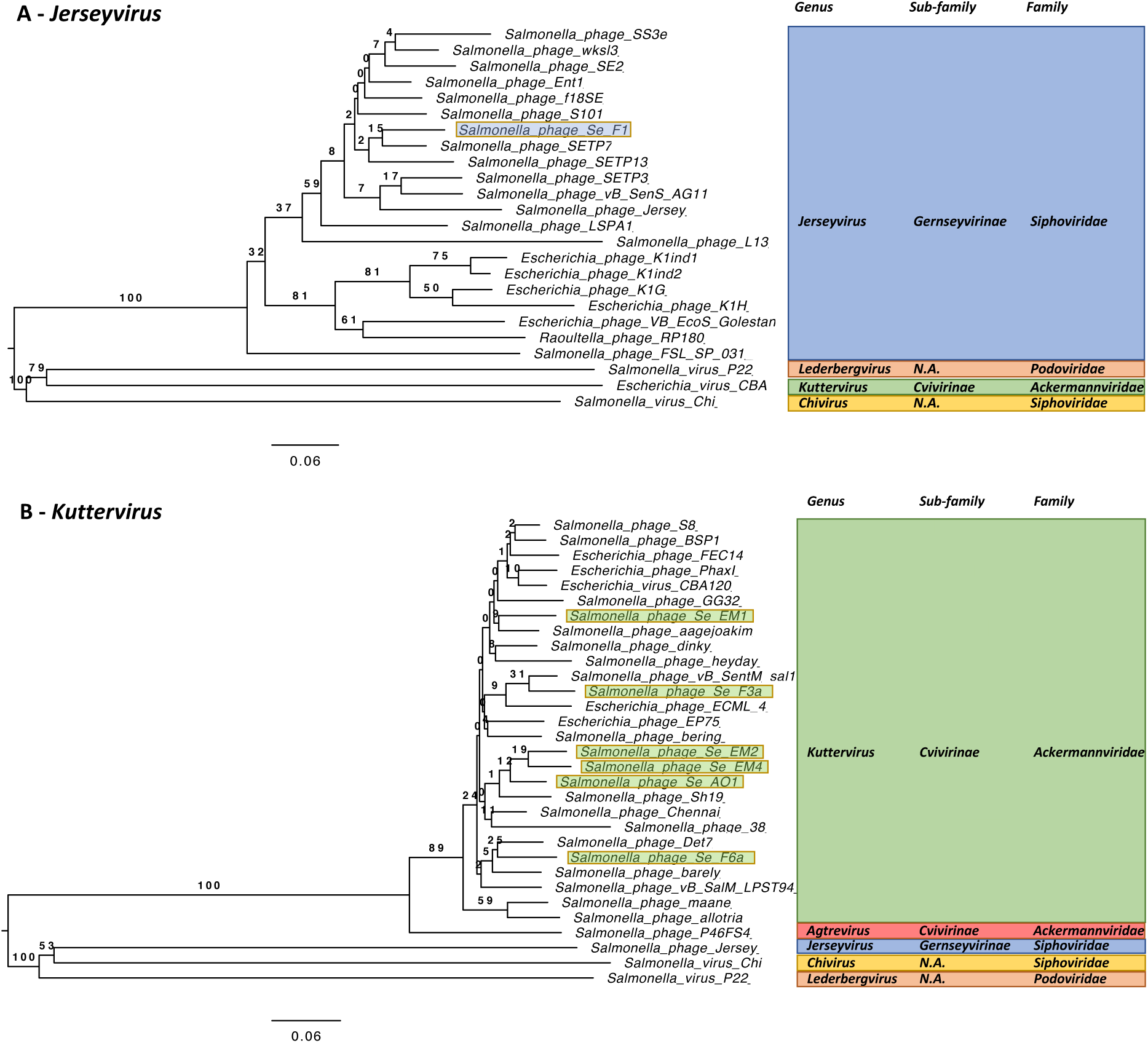
Taxonomic affiliation at the genus level based on genome wide, amino acid analysis by VICTOR. Panel A: *Jerseyvirus* genus. 20 *Jerseyvirus* from RefSeq database were included, as well as 1 *Kuttervirus* (*Escherichia* virus CBA120), 1 *Chivirus* (*Salmonella* phage Chi) and 1 *Lederbergvirus* (*Salmonella* phage P22) serving as outgroups. *Salmonella* phage Se_F1 is highlighted in light blue. Panel B: *Kuttervirus* genus. 21 *Kuttervirus* from RefSeq database were included, as well as 1 *Agtrevirus* (*Salmonella* phage P46FS4), 1 *Lederbergvirus* (*Salmonella* phage P22), 1 *Chivirus* (*Salmonella* phage Chi) and 1 *Jerseyvirus* (*Salmonella* phage Jersey) serving as outgroups. *Salmonella* phage Se_AO1, Se_EM1, Se_EM2, Se_EM4, Se_F3a, Se_F6a are highlighted in light green. All accession numbers are summarized in Supplementary Table 3.

Figure 4 unambiguously shows that *Salmonella* phage Se_F1 belongs to the *Jerseyvirus* genus and that *Salmonella* phages Se_AO1, Se_EM1, Se_EM2, Se_EM4, Se_F3a and Se_F6a belong to the *Kuttervirus* genus. Table 4 summarizes the final taxonomic affiliation of the 10 new phage species studied in this work.

**Table 4:** Taxonomic classification at the 10 newly isolated *Salmonella* phages included in this study. All phages in this table belong to the *Duplodnaviria* realm, *Heunggongvirae* kingdom, *Uroviricota* phylum, *Caudoviricetes* class, *Caudovirales* order. For illustration, in the “guest star” column are listed the most “famous” (or seeding) viruses belonging to the considered genus (accession numbers are listed in Supplementary Table 3).

| Phage species | Family | Sub-family | Genus | Guest star |
| --- | --- | --- | --- | --- |
| *Salmonella* phage Se_AO1 | *Ackermannviridae* | *Civivirinae* | *Kuttervirus* | *Escherichia* virus<br>CBA120 |
| *Salmonella* phage Se_EM1 |  |  |  |  |
| *Salmonella* phage Se_EM2 |  |  |  |  |
| *Salmonella* phage Se_EM4 |  |  |  |  |
| *Salmonella* phage Se_F3a |  |  |  |  |
| *Salmonella* phage Se_F6a |  |  |  |  |
| *Salmonella* phage Se_F1 | *Siphoviridae* | *Guernseyvirinae* | *Jerseyvirus* | *Salmonella* virus<br>Jersey |
| *Salmonella* phage Se_F2 | *Siphoviridae* | N.A. | *Chivirus* | *Salmonella* virus Chi |
| *Salmonella* phage Se_F6b |  |  |  |  |
| *Salmonella* phage Salfasec_13b | *Podoviridae* | N.A. | *Lederbergvirus* | *Salmonella* virus P22 |

This taxonomic classification allowed us to confirm the PhageTerm predictions for *Salmonella* phages Se_F2 and Se_F6b (*Chivirus* genus). The closest homologs in term of DNA sequences are *Salmonella* phage BPS1 and *Salmonella* phage Chi for *Salmonella* phages Se_F2 and Se_F6b, respectively (Table 2). In both cases, PhageTerm was able to predict a cos-type DNA packaging with the same single-stranded extremities than reported by Hendrix et *al.* by primer extension experiment for *Salmonella* phage Chi (5’-GGTGCGCAGAGC-3’) [60]. This gives credit to PhageTerm prediction for *Salmonella* phage Se_F6b despite the fact that the sequence coverage was below the recommended threshold (Table 3).

### IV.5 Functional annotation

To date, the most common and fastest method to tentatively ascribe a function to a newly predicted gene product is by comparing the ORF or gene product sequences with existing sequences deposited in various databases such as the nr/nt databases at NCBI. Sequence alignments are usually generated with now classical algorithms such as BLAST, T-Coffee, MUSCLE or Clustal Omega [61–64]. When the sequence homology is deemed sound, the annotation of the known gene function is then simply transferred to the newly predicted gene. This is of course very reliable when functions can be traced back to experimental validation. Synteny analysis is also an interesting approach to make educated guesses on the function of an unannotated gene by comparing similar, annotated genomic environments. For instance, Phagonaute [65] was developed for bacterial and archaeal viruses to compare the genetic context of a selected gene in parallel to the genetic context of its homologs in other sequenced phage genomes. Unfortunately, Phagonaute is not updated anymore. For proteins that finally end up with “unknown function” annotation, one has to keep a critical eye for several reasons: i) the function can truly be a novel one (the phage “dark-matter”), ii) the amino acid sequence is too divergent to detect homologies with proteins of known function using “simple” sequence alignment algorithms such as BlastP, or iii) the gene product does not exist (an artifact due to the ORF prediction algorithms). Functional annotation can be improved thanks to methods developed to detect remote homologies, classify protein in orthologous groups or fold primary sequences onto known secondary or tertiary structures. As noted by Chen et *al.* [66] these many different approaches can be combined to yield functional predictions as reliable as can be.

Here we chose to combine two complementary approaches for the functional annotation of our 10 phage genomes:

- Comparison of the predicted gene products with the newly published PHROG database comprising exclusively protein sequences from viruses infecting bacteria and archaea as well as their prophages. In PHROG, viral proteins are clustered in protein orthologs families called “phrog” built on remote homology detection and functionally annotated.
- Comparison of the predicted gene products with protein structures from the pdb70 database at the Protein Data Bank.

We thus first compared all predicted gene products for each phage genome with the PHROG database. Supplementary Excel file 3 recapitulates the best hit in PHROG for each predicted ORF as well as the comparison metrics. We then used the comparison metrics and the genomic environment to perform a manual curation of the predicted ORFs. In total, 59 ORFs were thus manually curated (highlighted in red in Supplementary Excel file 1 and in Supplementary Excel file 3), ranging from 1 curated ORF per genome (*Salmonella* phage Se_F2 and Se_F6b) to 10 curated ORFs per genome (*Salmonella* phage Se_AO1). 89.8% of these curated ORFs (53 ORFs) were predicted by Phanotate, suggesting that this gene caller designed for phage genomes tends to overpredict ORFs. For the remaining ORFs annotated unknown function with a weak affiliation to a phrog family we replace the phrog number by “singleton”, meaning that this ORF does not have (yet) any ortholog in the PHROG database. This can be either a false positive ORF or a truly novel function. The percentage of singletons varies from 8 to 13% depending on the genome (Table 6).

For gene products still annotated with an unknown function after comparison with the PHROG database, we tried to improve the annotation by comparison with the pdb70 database of the Protein Data Bank. The best hits for each gene product of each genome are listed in Supplementary Excel file 4. After manual inspection of the results and the comparison metrics, we could propose a functional annotation for 24 phrog families annotated unknown function. These propositions are summarized in Table 5.

**Table 5.**
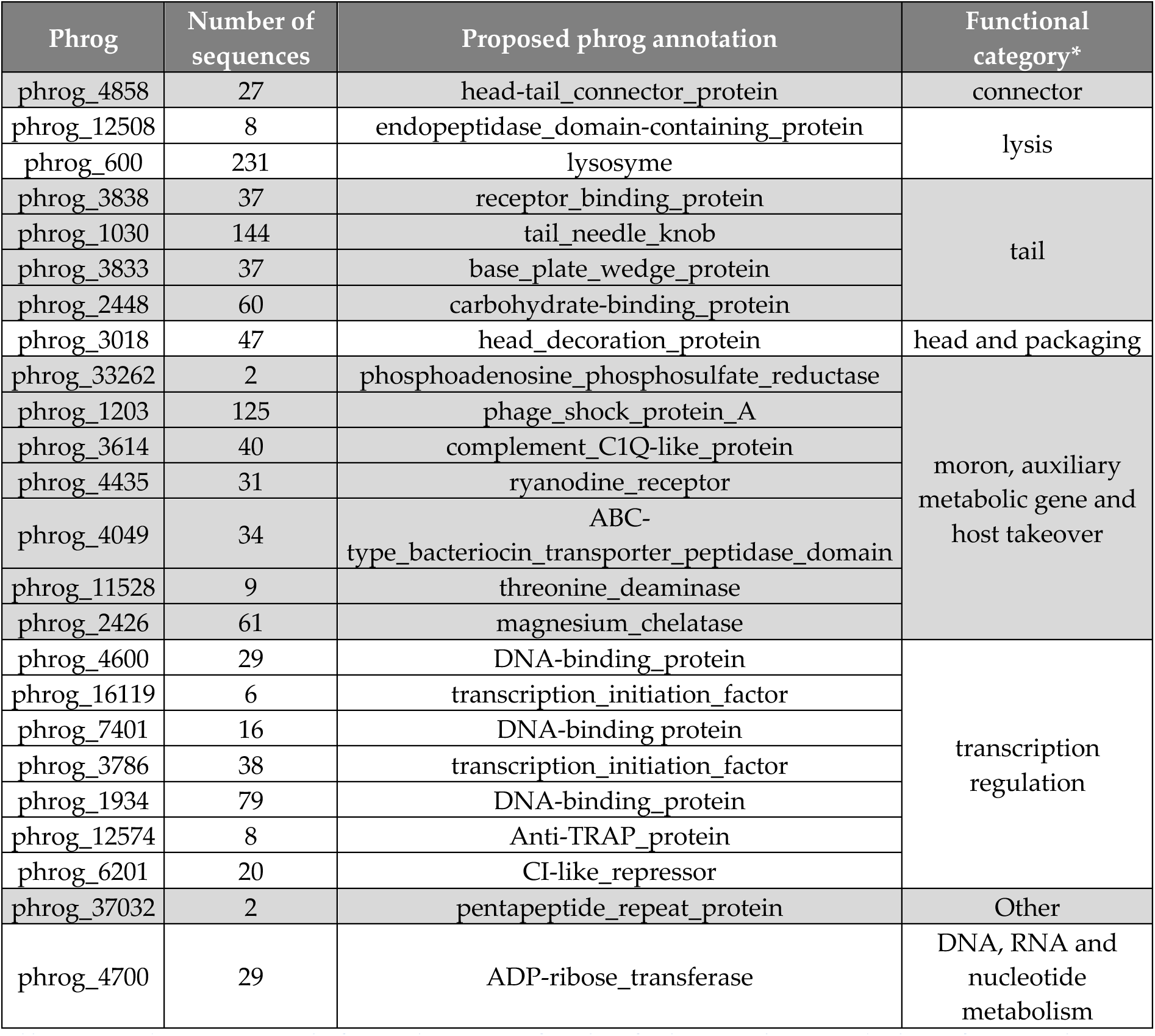
Suggested improvement to the functional annotation of 24 phrog families currently annotated unknown function in the PHROG database. *Functional categories defined in PHROG.

The proposed annotations cover 1,120 protein sequences included in the PHROG database, 352 of which are structural proteins and 196 participate to gene regulation according to our proposed annotations. Of note, phrog_6201 is predicted as a CI-like repressor, one of the usual markers of the temperate bacteriophages. This phrog contains 20 protein sequences from 20 distinct phages that are thus likely to be temperate phages. Of note, *Salmonella* phage Salfasec_13b Gp_057 protein belongs to the phrog_6201, hinting at a temperate lifestyle for this bacteriophage. This example illustrates as one can gain valuable functional insights on an entire set of bacteriophages just by improving functional annotation of a single phrog.

Table 6 summarizes the final syntaxic and functional annotation for each phage. Detailed functional annotations for each genome can be found in Supplementary Excel file 5.

**Table 6:** Summary of the final syntaxic and functional annotations of the 10 phage genomes. “Unknown function” represents the percentage of predicted ORFs annotated unknown function. “Singleton” represents the percentage of predicted ORFs that does not belong to any phrog family.

| Phage name | Length (bp) | Predicted ORFs | Predicted tRNAs | Unknown function | Singleton | Accession |
| --- | --- | --- | --- | --- | --- | --- |
| *Salmonella* phage Se_AO1 | 157543 | 233 | Asn, Gln, Met, Ser, Tyr, Val | 57.5% | 10.3% | Submitted |
| *Salmonella* phage Se_EM1 | 159312 | 236 | Asn, Met, Pro, Ser, Tyr | 57.2% | 11.0% | Submitted |
| *Salmonella* phage Se_EM2 | 156948 | 233 | Asn, Met, Pro, Ser, Tyr | 55.4% | 11.1% | Submitted |
| *Salmonella* phage Se_EM4 | 157810 | 231 | Asn, Met, Pro, Ser, Val | 55.0% | 10.8% | Submitted |
| *Salmonella* phage Se_F1 | 42784 | 74 | None | 37.8% | 8.1% | Submitted |
| *Salmonella* phage Se_F2 | 59084 | 86 | None | 51.2% | 12.8% | Submitted |
| *Salmonella* phage Se_F3a | 157219 | 236 | Asn, Ile, Met, Tyr | 55.5% | 10.2% | Submitted |
| *Salmonella* phage Se_F6a | 157240 | 233 | Asn, Ile, Pro, Ser, Tyr | 53.6% | 9.0% | Submitted |
| *Salmonella* phage Se_F6b | 59519 | 84 | None | 50.0% | 11.9% | Submitted |
| *Salmonella* phage Salfasec_13b | 40455 | 74 | Asn | 41.9% | 12.2% | Submitted |

The final GenBank files for submission were built using the PHROG annotation for each ORF; when pertinent the predicted functions derived from the PDB comparisons were added together with its corresponding PDB entry. As the PHROG database is not yet recognized by the International Nucleotide Sequence Database Collaboration (INSDC) it was not possible to use the “db_xref” flag for the PHROG annotations in the GenBank file. As a consequence, we reported the phrog number and the corresponding functional annotation for each gene as a “note” in the GenBank file. The 10 bacteriophage genomes were submitted to GenBank under the accession numbers listed in Table 6 and regrouped under the BioProject accession number PRJNA767534.

### IV.6 Phage life-style

Consensually, the development of phage therapy strategies relies on strictly virulent bacteriophages to formulate cocktails. The exclusion of temperate phages aims at avoiding lysogenization that 1) would not kill immediately the targeted cell, 2) could enable the acquisition of virulence genes or other genes increasing the fitness of the lysogens. In a somehow iconoclast view, Monteiro et *al.* [67] recently argued that temperate phage could be interesting as they are abundantly present in bacterial genomes, ready to be used and eventually engineered, although current regulations towards genetically modified organisms do not favor yet such approaches. Hence the prediction of phage lifestyle prior to therapeutical cocktail formulation still remains a prerequisite. We scanned each phage proteome for predicted functions related to the bacteriophage life-styles (virulent or temperate). When no integrase together with other functions generally associated with temperate phages such as a Recombination Directionality Factor (RDF, also called excisionase) or CI-like repressors were found, we hypothesized a “virulent” life style. Such was the case for all phages but *Salmonella* phage Salfasec_13b.

We believe that *Salmonella* phage Salfasec_13b (*Lederbergvirus* genus) is a temperate phage as it contains many functions associated with temperate phages such as an integrase of the tyrosine recombinase family (Gp_026, phrog_216), a CIII anti-termination protein (Gp_041, phrog_550), a CI-like repressor (Gp_057, phrog_6201) and a CII-like regulator (Gp_060, phrog_725). However, we could not detect a Recombination Directionality Factor (RDF or excisionase) required for prophage excision. RDFs are difficult to predict as they are small proteins sharing very few sequence homologies (66 aa for TorI, the RDF for the defective prophage KplEI in *E. coli* K12 [68, 69]). It is thus probable that Salfasec_13b RDF is located among the small ORFs of unknown function. RDF genes are commonly found in the vicinity of their cognate integrase genes. Of note, in *Salmonella* phage Salfasec_13b, the integrase gene (*gp_026*) is surrounded by a predicted ORFs encoding a small protein of unknown function (Gp_027, 56 aa) compatible with the size of an RDF.

*Salmonella* phage Se_F1 (*Jerseyvirus* genus) is an interesting case as it contains a predicted RDF (Gp_053, 76 aa, phrog_66) whose modeled structure we found matches very well with TorI (PDB 1Z4H), the RDF of the defective prophage KplE1 mentioned above. Nevertheless, no ORF coding for an integrase could be predicted and others key functions associated with temperate phages are missing. Same findings were made for the closest annotated sequence homolog found with Blastn in the nt database, *i.e., Salmonella* phage S101 (Table 2). The PHROG database includes 22 *Jerseyvirus* genomes, 20 of them coding for RDF belonging to the phrog_66, making it a marker of this genus although *Jerseyvirus* seem to be strictly virulent phages. *Salmonella* phage Se_F1 and its relatives might have been temperate phages that have lost their integrase gene or the RDF encoded in their genome has been acquired by horizontal gene transfer and may serve to induce resident prophage(s) in the host. The latter hypothesis is worth investigating as induction of resident prophages is likely to ensure an efficient lysis of the host.

### IV.7 Genetic mining of 10 newly isolated *Salmonella* phage genomes in the light of phage therapy

The primary aim of our isolation of new *Salmonella*-targeting phages being the development of phage therapy against *Salmonella* species, we were particularly interested in two major kinds of protein functions:

- Those that could give a selective advantage for host infection, successful virion productions and cell lysis (mostly features allowing evasion from cellular defenses).
- Those that could interfere with the formulation of a bacteriophage cocktail such as functions allowing lysogeny or super immunity against other phage infections.

We first reviewed these two categories of functions by bacteriophage genus before focusing on the prediction of anti-CRISPR proteins which belong to the first category of proteins likely to confer a selective advantage.

#### Jerseyvirus: Salmonella *phage Se_F1*

We identified a DarB-like Type-I RM anti-restriction protein (Gp_039, phrog_10089). In bacteriophage P1 infecting *E. coli*, several anti-restriction proteins among which DarB are packaged during virion assembly and released upon infection into the host cytoplasm, ready to act against host RM systems [70]. DarB alone is required for protection against type-I EcoB and EcoK restriction. The Gp_039 annotation (DarB-like antirestriction) is sound because this protein belongs to phrog_10089 predicted to be highly similar (probability 100%, Evalue 3E-210) to phrog_1685 that does contain Bacteriophage P1 DarB. It thus seems that *Salmonella* phage Se_F1 encodes at least a DarB-like anti-restriction protein. Of note, *Salmonella* phage Se_F1 genome contains four EcoB and four EcoK sites.

Gp_023 is predicted as an anti-repressor of the Ant type (phrog_130). In *Salmonella* prophages such as Gifsy-1 and Gifsy-2, prophage induction is repressed by the binding of the repressor Rep on the promoter upstream genes allowing prophage induction [71]. The production of Ant derepresses these genes by titrating the Rep protein, thus triggering prophage(s) induction and eventually cell lysis. Gp_023 expression is then susceptible to induce resident prophage(s) in the infected host and may then help for host takeover. Together with the RDF mentioned previously, this is the second protein that is indicative of a temperate lifestyle, although *Salmonella* phage Se_F1 lacks the other functions required for lysogenization.

Gp_027 is predicted as an immunity-to-superinfection protein (phrog_1039). In *E. coli* phage T4, the immunity protein Imm blocks the injection of T4 DNA, preventing superinfection of phages belonging to the same family. Bacteriophage T4 Imm belongs to the same phrog as Gp_027, strengthening the proposed annotation for Gp_027. The production of an Imm-like protein in cells infected by *Salmonella* phage Se_F1 may thus lead to the exclusion of other phages in a cocktail that belong to the same family. In such a situation, using two or more phages of the same family in the same cocktail is useless.

*Salmonella* phage Se_F1 seems to harbor an extensive arsenal for host lysis with two Rz-like spanins (Gp_009 and Gp_010, phrog_2536 and phrog_5672, respectively), one endopeptidase predicted from structural comparison with the pdb70 (Gp_034, phrog_12508 currently annotated “unknow function” in the PHROG database), two holins (Gp_063 and Gp_064, phrog_2359 and phrog_2508, respectively) and one endolysin (Gp_065, phrog_33083).

#### Chivirus: Salmonella phages Se_F2 and Se_F6b

*Salmonella* phage Chi and other Chi-like viruses are flagellotropic phages requiring a motile flagellum as its primary receptors; it is then very likely that *Salmonella* phages Se_F2 and Se_F6b share the same requirement. Both *Salmonella* phages Se_F2 and Se_F6b phages contain 3 gene products with predicted carbohydrate-binding domains we predicted from structural comparison with the pdb70 (phrog_2448 currently annotated “unknown function” in the PHROG database) encoded on the same strand downstream of the tail genes module, suggesting that these proteins may be part of the tail and the host recognition apparatus. These genes encoding carbohydrate-binding domains are shared among 36 other *Chivirus* present in the PHROG database. Sugars are components of the LPS and it has also been shown that *S. typhimurium* flagella contain at least 16 different sugars [72]. The flagellotropic *Salmonella* phage Chi and its relatives may use these domains to recognize and bind the bacterial flagellum and its sugar moieties. It has been reported that *Salmonella* phage Chi can infect both *Salmonella spp* and *E. coli* hosts and thus has been proposed as a good candidate for the development of phage-based applications (pathogens detection, remediation, phage therapy) [73]. Inclusion in a therapeutical cocktail of bacteriophages such as *Chivirus* that may target other receptor than the LPS would be a good option to avoid cross-resistance.

Both phages encode a predicted DNA methyltransferase (phrog_111) homologous to the DNA N-6-adenine methyltransferase according to PFAM prediction (PF05869) attached to this phrog. This enzyme may serve to modify phage DNA during replication, allowing the phages to escape host restriction-modification systems.

#### Lederbergvirus: Salmonella *phage Salfasec_13b*

As discussed above, *Salmonella* phage Salfasec_13b is a temperate phage and a *Lederbergvirus* (as the archetypal *Salmonella* phage P22). This phage genome contains all the genes found in P22 that are important to promote lysogeny or confer a selective advantage to the host including *gtrA* (Gp_022, phrog_2224), *gtrB* (Gp_021, phrog_34335) and *gtrC* (Gp_019, phrog_4829). Together the GtrABC complex modifies the host O-antigen at the host cell surface [74]. Changing the LPS prevents infection by other phages using it as a primary receptor. In the light of phage therapy this is not a desired outcome.

#### Kuttervirus: Salmonella phages Se_AO1, Se_EM1, Se_EM2, Se_EM4, Se_F3a, Se_F6a

We first pinpoint protein functions that seem to be a hallmark of the *Kuttervirus* genus as they are found both in our 6 *Kuttervirus* phage genomes but also in almost all the 15 *Kuttervirus* genomes included in the PHROG database.

An interesting feature is the gene product belonging to phrog_1510 (superinfection exclusion) that includes the Gp17 protein of *Salmonella* phage P22. In the classical literature, Gp17 is described as a protein necessary for P22 to counteract a superinfection exclusion system encoded in the Fels-2 prophage found in many *S. enterica Typhimurium* strains [75]. A sensible hypothesis is that *Kuttervirus* phages have somehow acquired a P22 *gp17-like* gene that allows them to successfully infect and propagate in lysogenic *Salmonella* strains harboring Fels-2-like superinfection exclusion systems. This is definitively an interesting feature for a phage cocktail as *Salmonella spp* harbor many and diverse prophages [76].

Another hallmark is phrog_600 (unknown function) that we could predict as a lysozyme-like protein by comparison with the pdb70. This adds to the predicted Rz-spanin and endolysin in the arsenal of proteins ensuring host lysis.

The RIIA (phrog_3661, RIIA lysis inhibition) and RIIB (phrog_3803, RIIB lysis inhibition) are also encoded in most *Kuttervirus* phages. The phage proteins RIIA and RIIB enable bacteriophage T4 to resist to the Rex restriction system encoded by *E. coli* Lambda lysogens. Rex is an abortive infection defense system and relies on the action of two proteins, RexA and RexB [77]. However, the molecular mechanisms behind Rex still remains elusive. RIIA and RIIB could confer an advantage to *Kuttervirus* phages when infecting lysogens encoding a Rex-like system.

We were intrigued by the presence in all our six *Kuttervirus* genome of a predicted second terminase small subunit (phrog_800) situated far away from the traditional pair of consecutive genes encoding the small and the large terminase subunits. To the best of our knowledge, this is unheard of in phage genomes. A closer inspection of phrog_800 revealed a probable misannotation. Indeed, by checking the list of PDB hits (Supplementary Excel file 4) we found out that phrog_800 proteins are predicted to be deoxynucleoside kinase proteins. In our genomes, phrog_800-encoding gene is followed by genes encoding a thymidiate synthase, a deoxynucleoside monophosphate kinase and finally a nucleoside triphosphate pyrophosphohydrolase. This genomic environment strongly suggests a gene cluster involved in DNA metabolism and strengthens our proposed new annotation for phrog_800 as a deoxynucleoside_kinase.

Finally, an interesting feature was found only in *Salmonella* phage Se_F3a. Our comparison with the pdb70 strongly suggests that Gp_125 (phrog_1203, unknown function) is homologous to PspA, the Phage Shock Protein A. Four of the 15 *Kuttervirus* phage genomes included in the PHROG database harbor such a gene. *pspA* belongs to the *psp* operon found in some bacterial genomes, noticeably in *E. coli* and *Salmonella* species. This system prevents the dissipation of the proton-motive force (PMF) triggered by various stresses, including the infection by filamentous phages or the mislocalization of secretins [78]. PspA plays a role in the regulation of the phage-shock-response *psp* operon but also associates with the inner membrane to prevent the loss of PMF, although the molecular mechanisms still remain unclear. Several Abi systems such as Rex in Lambda lysogens mentioned above or AbiZ in *Lactococcus lactis* [13] lead to cell death or cellular growth arrest through membrane permeabilization and dissipation of the PMF or loss of ATP. A reasonable hypothesis could be that the phage-encoded PspA acts as a novel anti-Abi system, preventing cell suicide. This hypothesis will be worth further investigations. This feature singles out *Salmonella* phage Se_F3a from the other *Kuttervirus* phages isolated in this study, with a putative additional ant-host defense to its arsenal.

#### Prediction of anti-CRISPR proteins (Acr)

We performed Acr encoding genes prediction with AcrHub. Among all the Acr predicted in our 10 phage genomes, we retained the most probable predictions summarized in Table 7.

**Table 7:** Predicted Acr proteins in the 10 phage genomes studied. This table is a simplified version of Erreur ! Source du renvoi introuvable.. Scores for two different Acr predictors, PaCRISPR, AcRanker, are indicated:. Column 7: The closest homologs in the AcrHub database are displayed^#^, as well as the inhibited CRISPR types^&^. *PaCRISPR score below 0.5 but this predicted Acr belongs to the phrog_519 that found in the other five *Kuttervirus* genomes with reliable scores.

| Phage | Genus | phrog | Protein | Size (aa) | PaCRISPR score | AcRanker score | AcrHub ID<sup>‡</sup> | CRISPR Type<sup>&</sup> |
| --- | --- | --- | --- | --- | --- | --- | --- | --- |
| *Salmonella* phage Se_AO1 | *Kuttervirus* | 4097 | Gp_007 | 56 | 0.62 | -0.75 | Acr00210 | II-A |
|  |  | 519 | Gp_015 | 71 | 0.54 | -2.08 | Acr00037 | I-D |
|  |  | 5934 | Gp_049 | 105 | 0.69 | -4.64 | Acr00193 | II-A |
|  |  | 3594 | Gp_088 | 62 | 0.58 | -3.57 | Acr00054 | I-D |
| *Salmonella* phage Se_EM1 | *Kuttervirus* | 4097 | Gp_008 | 56 | 0.62 | -0.75 | Acr00210 | II-A |
|  |  | 3594 | Gp_080 | 62 | 0.55 | -2.90 | Acr00212 | II-A |
| *Salmonella* phage Se_EM2 | *Kuttervirus* | 4097 | Gp_008 | 56 | 0.62 | -0.75 | Acr00210 | II-A |
|  |  | 519 | GP_016 | 71 | 0.54 | -2.08 | Acr00037 | I-D |
| *Salmonella* phage Se_EM4 | *Kuttervirus* | 4097 | Gp_007 | 56 | 0.62 | -0.75 | Acr00210 | II-A |
|  |  | 519 | Gp_015 | 71 | 0.54 | -2.08 | Acr00037 | I-D |
| *Salmonella* phage Se_F3a | *Kuttervirus* | 4097 | Gp_008 | 56 | 0.62 | -0.74 | Acr00210 | II-A |
|  |  | 519 | Gp_014 | 78 | 0.48* | -1.17 | Acr00037 | I-D |
|  |  | 3594 | Gp_082 | 62 | 0.54 | -2.77 | Acr00212 | II-A |
| *Salmonella* phage Se_F6a | *Kuttervirus* | 4097 | Gp_007 | 56 | 0.62 | -0.74 | Acr00210 | II-A |
|  |  | 5934 | Gp_042 | 110 | 0.68 | -3.85 | Acr00211 | II-A |
| *Salmonella* phage Se_F1 | *Jerseyvirus* | 13851 | Gp_038 | 73 | 0.58 | -4.11 | Acr00209 | II-A |
|  |  | 18511 | Gp_057 | 61 | 0.53 | -1.96 | Acr00311 | V-A |
|  |  | 6094 | Gp_059 | 173 | 0.83 | -3.57 | Acr00061 | I-E |
|  |  | 2180 | Gp_068 | 61 | 0.53 | -4.87 | Acr00128 | I-F |
|  |  | 1402 | Gp_072 | 69 | 0.64 | -3.92 | Acr00070 | I-F |
|  |  | Singleton | Gp_073 | 62 | 0.70 | -4.91 | Acr00264 | II-C |
| *Salmonella* phage Se_F2 | *Chivirus* | 7965 | Gp_051 | 120 | 0.84 | -4.46 | Acr00254 | II-C |
|  |  | 7734 | Gp_052 | 111 | 0.54 | -4.86 | Acr00029 | I-D |
|  |  | 4445 | Gp_057 | 85 | 0.77 | -5.17 | Acr00159 | II-A |
| *Salmonella* phage Se_F6b | *Chivirus* | 7965 | Gp_051 | 120 | 0.84 | -4.21 | Acr00275 | II-C |
|  |  | 7734 | Gp_052 | 111 | 0.53 | -4.80 | Acr00029 | I-D |
|  |  | 4445 | Gp_058 | 86 | 0.69 | -4.94 | Acr00198 | II-A |
| *Salmonella* phage Salfasec_13b | *Lederbergvirus* | 11224 | Gp_044 | 41 | 0.65 | -4.56 | Acr00064 | I-E |

In all phage genomes studied here, we could retain at least one Acr encoding gene prediction. *Salmonella* phage Se_F1 harbors the highest number of predicted Acr (*n* = 6). These predictions are generally consistent for bacteriophages belonging to the same genus. For *Kuttervirus* genomes, phrog_4097 is found in all six genomes studied and phrog_519 in five over six genomes. For *Chivirus* genomes, *Salmonella* phages Se_F2 and Se_F6b genomes code for the same three predicted Acr.

Interestingly, the transposable *Pseudomonas aeruginosa* Mu-like phage genomes contain an *acr* locus comprising two to three genes, located among the genes involved in the phage head assembly [52]. In these phages, the *acr* locus is bracketed by genes involved in phage head morphogenesis, usually upstream a protease/scaffold gene. The lower panel in Figure 5 describes the genetic context of AcrIE3, a type-IE Acr protein from *P. aeruginosa* phage DMS3 that has been functionally investigated [79]. In our six *Kuttervirus* phages we found one small protein (56 aa) belonging to phrog_4097 (unknown function) predicted with good scores by both PaCRISPR and AcRanker as a type II-A anti-CRISPR protein. The gene encoding this protein in the *Kuttervirus* genomes is situated between the portal protein and a head scaffolding protein encoding genes, reminiscent of the genetic context of AcrIE3 (Figure 5, upper panel).

**Figure 5:**
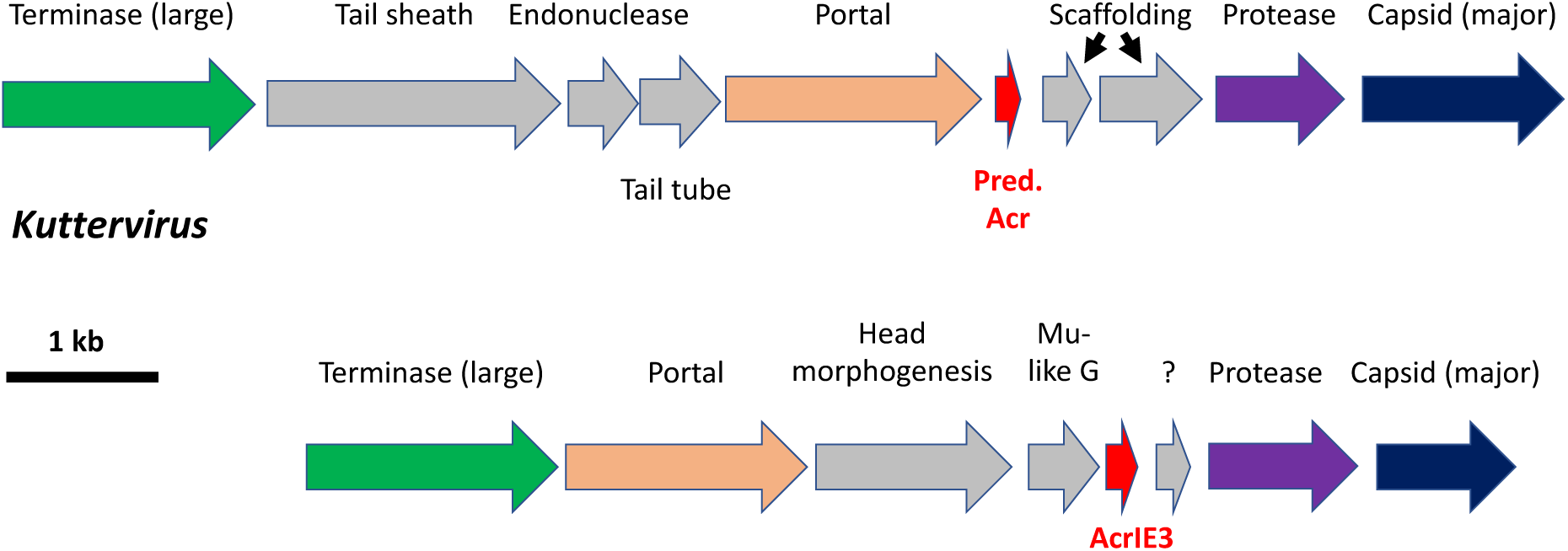
Comparison of the genomic surroundings of a *bona fide* Acr-encoding genes (*acrIF2*) in Mu-like *Pseudomonas aeruginosa* phage DMS3 (accession NC_008717) and the predicted Acr-encoding ORF in our 6 *Kuttervirus*. For clarity, genes are drawn to scale but not the intergenic regions.

#### Mu-like *P. aeruginosa* phage DMS3

One can also find a similar a gene coding for a phrog_4097 protein in the same genomic context in 14 of the 15 Kuttervirus genomes found in the PHROG database, making this predicted Acr another hallmark of *Kuttervirus* genomes. This predicted Acr is thus a serious candidate for anti-CRISPR functional studies and an interesting feature for therapeutical cocktails.

*Salmonella* phage Se_F1 Gp_057 exhibits high scores for both PaCRISPR and AcRanker predictions. Although we could not link *gp_057* genomic context to known contexts of experimentally validated Acr, we believe Gp_057 is also a serious candidate for anti-CRISPR functional studies.

## V. CONCLUSION

In this study, we present a genomic description of 10 distinct new phages isolated in wastewater and fresh water pounds in the Sevilla area in Spain and infecting *S. enterica* serovar *Typhimurium* strain ATCC 14028S. Based on genome-wide analyses independent from prior knowledge on functional annotation, we could ascribe taxonomic classification down to the genus level. These phages all belong to the *Caudovirales* (dsDNA tailed bacteriophages) and to four different genera (Figure 3 and Figure 4). *Kuttervirus* phages are overrepresented with six species identified. Only *Salmonella* phage Salfasec_13b, belonging to the *Lederbergvirus* genus, was predicted as a temperate phage.

We wish to highlight in our study two important methodological points. The first one is that by combining the results of four different gene callers – included Phanotate especially designed for phage genomes (but that nevertheless tends to over-predict ORFs), we believe we could predict as many ORFs as it is reasonably possible. However, we also showed that manual inspection and curation is still needed to get rid of obvious false positive predictions. The second one concerns the improvement of functional annotation with PHROG, a database dedicated to viral proteins at large (bacterial and archaeal viruses and their prophages). We believe this is a promising approach for functional annotation as it relies on viral protein orthologs clustering based on remote homology detection. PHROG capitalizes on past work of many research teams for decades that led to the experimental validations of many functions and PHROG also benefits from manual annotation of various experts in the field (we could propose in this study a functional annotation to 24 phrog clusters previously annotated with unknown function).

We mined our 10 annotated genomes for relevant functions in the context of future biocontrol application of *Salmonella spp*. We could thus identify phage-encoded proteins targeting bacterial anti-phage defenses such as the Abortive infection, Restriction-Modification and CRISPR-Cas systems. Our study illustrates also that in otherwise very similar genomes such as our six *Kuttervirus* (ANI values between 0.94 and 0.98) we could identify subtle variations in gene equipment that can single out one phage from its closest relatives. Indeed, we predicted that *Salmonella* phage Se_F3a genome alone codes for a new anti-Abi system based on a phage-encoded Phage Shock Protein A ortholog, potentially conferring *Salmonella* phage Se_F3a a selective advantage over the other five viruses of the same *Kuttervirus* genus. This example emphasizes the need to extensively mine bacteriophage genomes in order to tailor therapeutical cocktails matching the targeted pathogen. Conversely one has to obtain as many genomic information as possible about the pathogen itself in order to identify its anti-phage defenses and select appropriate phages to formulate a therapeutical cocktail that can overcome these cellular defenses. The recent publication of Tesson et *al.* comes handy in that respect with the description of DefenseFinder, a bioinformatic tool especially designed to systematically predict anti-phage systems in prokaryotic genomes [80].

Coevolution of bacteria and their viruses for ages beyond count is often described as an arm race that gifted both protagonists with highly diverse molecular weapons and shields. To go beyond empiric approaches and rationally design efficient phage-based biocontrol of bacteria we need to adopt a holistic approach of the bacteriophage / host binomen. In that respect, genomic and functional analyses of both partners are crucial to meet the challenges already existing to tackle bacterial infections.

## VI. ACKNOWLEDGMENTS

We thank Artemis Kosta and Hugo Le Guenno from the Microscopy facility at the Institut de Microbiologie de la Méditerranée (IMM, FR 3479, CNRS – Aix-Marseille Université, Marseille, France) for TEM grids preparation and image acquisition. We are very thankful to Dr. Marie-Agnès Petit (INRAE, Jouy-en-Josas, France) for fruitful discussions on functional prediction and phage genome annotation, discussions we had personally but also during dedicated workshops organized by the French bacteriophage network (www.phages.fr). We thank Pr. François Enault and Eric Olo Ndela (LMGE, Université Clermont Auvergne, Clermont-Ferrand, France) for their help with the PHROG database. We are very grateful to the “Phage Cycle and Bacterial Metabolism” team members at the Laboratoire de Chimie Bactérienne (UMR 7283, CNRS – Aix-Marseille Université, Marseille, France) for their help and support. We are also grateful to Modesto Carballo, Laura Navarro and Cristina Reyes from the Servicio de Biología, CITIUS, Universidad de Sevilla, for help with experiments performed at the facility.

## VIII. SUPPLEMENTARY TABLES

**Supplementary Table 1:** Summary of the 18 meaningful contigs obtained after SPAdes *de novo* assembly of the trimmed sequencing paired reads. *3 different sequencing runs were performed at different time and in different conditions (*e.g.*, number of genomes loaded onto the Illumina flow cell), accounting in part for the differences observed in sequence coverage.

| BioSample | Contig name | Contig size (bp) | Coverage* |
| --- | --- | --- | --- |
| Se_AO1 | Salmonella phage Se_AO1 | 157620 | 58 |
| Se_EM1 | Salmonella phage Se_EM1 | 159389 | 42 |
| Se_EM2 | Salmonella phage Se_EM2 | 157025 | 56 |
| **Se_EM3** | *Salmonella* phage Se_EM3 | 157176 | 87 |
| **Se_EM4** | *Salmonella* phage Se_EM4 | 157887 | 116 |
| **Se_ML1** | *Salmonella* phage Se_ML1 | 157296 | 67 |
| **Salfasec_9** | *Salmonella* phage Salfasec_9 | 43186 | 504 |
| **Salfasec_10** | *Salmonella* phage Salfasec_10 | 43187 | 269 |
| **Salfasec_11** | *Salmonella* phage Salfasec_11a | 59161 | 49 |
|  | *Salmonella* phage Salfasec_11b | 42744 | 870 |
| **Salfasec_13** | *Salmonella* phage Salfasec_13a | 157296 | 79 |
|  | *Salmonella* phage Salfasec_13b | 40532 | 259 |
| **Se_F1** | *Salmonella* phage Se_F1 | 42861 | 796 |
| **Se_F2** | *Salmonella* phage Se_F2 | 59161 | 685 |
| **Se_F3** | *Salmonella* phage Se_F3a | 157296 | 362 |
|  | *Salmonella* phage Se_F3b | 59161 | 15 |
| **Se_F6** | *Salmonella* phage Se_F6a | 157317 | 354 |
|  | *Salmonella* phage Se_F6b | 59596 | 15 |

**Supplementary Table 2:** Summary of the ORFs predictions for each phage genome. Column 2: total number of ORFs predicted with four gene callers. Column 3: percentage of ORFs predicted by all four gene callers. Column 4: percentage of ORFs only predicted by Phanotate.

| Phage name | ORFs<br>(total) | Predicted<br>by all | Only Predicted<br>by Phanotate |
| --- | --- | --- | --- |
| *Salmonella* phage Se_AO1 | 243 | 76.5% | 14.0% |
| *Salmonella* phage Se_EM1 | 242 | 73.5% | 14.5% |
| *Salmonella* phage Se_EM2 | 239 | 73.6% | 15.1% |
| *Salmonella* phage Se_EM4 | 240 | 73.3% | 14.6% |
| *Salmonella* phage Se_F1 | 76 | 72.2% | 11.8% |
| *Salmonella* phage Se_F2 | 87 | 74.7% | 14.9% |
| *Salmonella* phage Se_F3a | 243 | 70.8% | 15.2% |
| *Salmonella* phage Se_F6a | 242 | 73.5% | 12.8% |
| *Salmonella* phage Se_F6b | 85 | 75.3% | 14.1% |
| *Salmonella* phage Salfasec_13b | 82 | 61% | 18.3% |

**Supplementary Table 3:** Phage genome accession numbers used for VICTOR analyses. *According to the current ICTV taxonomic classification.

| Sequence name | Accession number | Genus* |
| --- | --- | --- |
| *Escherichia* phage K1G | NC_027993 | *Jerseyvirus* |
| *Escherichia* K1H | NC_027994 | *Jerseyvirus* |
| *Escherichia* phage Kind1 | NC_041897 | *Jerseyvirus* |
| *Escherichia* phage Kind2 | NC_041898 | *Jerseyvirus* |
| *Escherichia* phage VB_EcoS-Golestan | NC_042084 | *Jerseyvirus* |
| *Raoultella* phage RP180 | NC_048181 | *Jerseyvirus* |
| *Salmonella* phage f18SE | NC_028698 | *Jerseyvirus* |
| *Salmonella* phage Jersey | NC_021777 | *Jerseyvirus* |
| *Salmonella* phage L13 | NC_021317 | *Jerseyvirus* |
| *Salmonella* phage LSPA1 | NC_026017 | *Jerseyvirus* |
| *Salmonella* phage S101 | MH370359 | *Jerseyvirus* |
| *Salmonella* phage SE2 | JQ007353 | *Jerseyvirus* |
| *Salmonella* phage SETP3 | NC_009232 | *Jerseyvirus* |
| *Salmonella* phage SETP7 | NC_022754 | *Jerseyvirus* |
| *Salmonella* phage SETP13 | NC_022752 | *Jerseyvirus* |
| *Salmonella* phage SS3e | NC_006940 | *Jerseyvirus* |
| *Salmonella* phage vB_SenS_AG11 | NC_041991 | *Jerseyvirus* |
| *Salmonella* phage vB_SenS_Ent1 | NC_019539 | *Jerseyvirus* |
| *Salmonella* phage wksl3 | NC_041992 | *Jerseyvirus* |
| *Salmonella* phage FSL_SP-031 | NC_021775 | *Jerseyvirus* |
| *Escherichia* phage ECML-4 | NC_025446 | *Kuttervirus* |
| *Escherichia* phage EP75 | NC_049433 | *Kuttervirus* |
| *Escherichia* phage FEC14 | NC_049427 | *Kuttervirus* |
| *Escherichia* phage PhaxI | NC_019452 | *Kuttervirus* |
| *Escherichia* virus CBA120 | JN593240 | *Kuttervirus* |
| *Salmonella* phage 38 | NC_029042 | *Kuttervirus* |
| *Salmonella* phage aagejoakim | NC_049503 | *Kuttervirus* |
| *Salmonella* phage Chennai | MN953776 | *Kuttervirus* |
| *Salmonella* phage Det7 | NC_027119 | *Kuttervirus* |
| *Salmonella* phage dinky | NC_049504 | *Kuttervirus* |
| *Salmonella* phage GG32 | NC_031045 | *Kuttervirus* |
| *Salmonella* phage heyday | NC_049500 | *Kuttervirus* |
| *Salmonella* phage maane | NC_049508 | *Kuttervirus* |
| *Salmonella* phage PhiSH19 | NC_019530 | *Kuttervirus* |
| *Salmonella* phage S8 | KY630163 | *Kuttervirus* |
| *Salmonella* phage vB_SalM-LPST94 | MH523359 | *Kuttervirus* |
| *Salmonella* phage vB-SentM_sal1 | MT499896 | *Kuttervirus* |
| *Salmonella* virus allotria | NC_0495501 | *Kuttervirus* |
| *Salmonella* virus barely | NC_049505 | *Kuttervirus* |
| *Salmonella* virus bering | NC_049502 | *Kuttervirus* |
| *Salmonella* virus BSP101 | NC_049502 | *Kuttervirus* |
| *Salmonella* phage Chi | NC_025442 | *Chivirus* |
| *Salmonella* phage P22 | NC_002371 | *Lederbergvirus* |
| *Salmonella* phage P46FS6 | NC_049509 | *Agtrevirus* |

## IX. SUPPLEMENTARY EXCEL FILES

Supplementary Excel file 1: ORFs predicted by AMIGene, MetaGeneAnnotator, Phanotate and Prodigal for each genome. True: ORF predicted by the considered gene caller. Empty cell: ORF not predicted by the considered gene caller. In red: manually curated ORFs. In green: ORFs later annotated as phrog singleton of unknown function.

Supplementary Excel file 2: Viral clusters (VC) defined by vContact2.

Supplementary Excel file 3: Primary assignment of each predicted gene product from each genome to a phrog family. In red: ORFs that will be curated for downstream analyses.

Supplementary Excel file 4: PDB hits.

Supplementary Excel file 5: Functional annotation for each genome after manual curation. In green: ORFs annotated with unknown function before comparison with the pdb70.

